# Contribution of fungal microbiome to intestinal physiology, early-life immune development and mucosal inflammation in mice

**DOI:** 10.1101/819979

**Authors:** Erik van Tilburg Bernardes, Veronika Kuchařová Pettersen, Mackenzie W. Gutierrez, Isabelle Laforest-Lapointe, Nicholas G. Jendzjowsky, Jean-Baptiste Cavin, Fernando A. Vicentini, Catherine M. Keenan, Hena R. Ramay, Jumana Samara, Wallace K. MacNaughton, Richard J. A. Wilson, Margaret M. Kelly, Kathy D. McCoy, Keith A. Sharkey, Marie-Claire Arrieta

## Abstract

Gut microbiomes make major contributions to the physiological and immunological development of the host, but the relative importance of their bacterial and fungal components, and how they interact, remain largely unknown. We applied carefully controlled experiments in gnotobiotic mice colonized with defined communities of bacteria, fungi, or both to differentiate the direct role of fungi on microbiome assembly, host development, and susceptibility to colitis and airway inflammation. Our results revealed that fungal colonization alone was insufficient to promote the intestinal anatomic and physiological changes seen in mice colonized by bacteria, and had limited impact on the fecal metabolome. However, fungal colonization promoted major shifts in bacterial microbiome ecology, and had an independent effect on the innate and adaptive immune development in young mice. Fungi further exacerbated some aspects of the inflammatory effects of the bacterial community during OVA-induced airway inflammation by promoting macrophage infiltration in the airway. Our results demonstrate a dominant ecological and physiological role of bacteria in gut microbiomes, but highlight fungi as an ecological factor shaping the assembly of the bacterial community and a direct capacity to impact immune system and modulate disease susceptibility. These findings demonstrate that studies focused on bacteria alone provide an incomplete portrayal on microbiome ecology and functionality, and prompt for the inclusion of fungi in human microbiome studies.

## Introduction

The mammalian intestinal microbiome constitutes a complex community of prokaryotic and eukaryotic microorganisms across high-level clades of the tree of life^1–3^. Structural and metabolic signals derived from this microbial ecosystem engage in crosstalk that helps define the trajectory towards host health or disease, especially early in life^4–7^. The early stages of gut microbiome establishment is more dynamic when compared to adult microbiomes, and environmental factors can influence the composition and diversity of this community, as well as the trajectory of its assembly^7^. Disruptions of microbiome assembly during early life can lead to dysregulation of host physiology and increased susceptibility to immune, metabolic, neurological and psychiatric diseases^8, 9^. As a result, the study of the causes and health consequences of microbial alterations has received significant attention. However, most of what is known about the ecology of the gut microbiome and the patterns associated with human disease are limited to bacteria^10^.

Fungi are an integral part of a wide variety of microbial environments^11^, including the human gut^12–14^. Gut bacterial and fungal communities inhabit similar intestinal habitats^15^, and co-interact throughout the early stages of colonization^7^. Alterations in the gut fungal community (mycobiome) have also been implicated in human disease^16, 17^. Work from Sokol *et al.* revealed mycobiome alterations in inflammatory bowel disease (IBD) patients experiencing a flare compared to a healthy cohort or IBD patients in remission^18^. These alterations included an increased fungi/bacteria diversity ratio and an increased abundance of *Candida albicans*, suggestive of fungal overgrowth by this common yeast^18^. As well, another common yeast, *Malassezia restricta*, was present in the majority of patients carrying the IBD risk allele *CARD9*, a molecule involved in fungal innate immunity^19^. Experimental colonization of mice with *M. restricta* resulted in exacerbated Dextran Sodium Sulfate (DSS)-induced colitis, characterized by CARD9-dependent Th1 and Th17 inflammation^19^.

Early-life gut dysbiosis has also been linked to atopic asthma. We had previously shown that early-life bacterial communities are altered in Canadian infants that later developed atopic-wheeze^20^. Fujimura *et al.* extended our findings and revealed an expansion of *Candida sp.* and *Rhodotorula sp.* in stool samples from US infants that later developed atopy^13^. Similarly, we detected mycobiome alterations in stool of rural Ecuadorian infants that developed atopic-wheeze at 5 years^12^. Differences in the fungal community were more strongly associated with asthma risk than bacterial dysbiosis in the Ecuadorian study, in which we detected an overrepresentation of total fungal sequences and an expansion of the yeast *Pichia kudriavzevii* in children who later developed symptoms^12^. While these studies revealed interesting associations between early-life mycobiome alterations and infant atopy and asthma, the causal role of fungi in altering early-life microbiome assembly and asthma pathology has not been established to date.

A limited number of studies suggest an immunomodulatory role of commensal gut fungi. Antibiotic-induced overgrowth of *C. albicans* or *C. parapilopsis* in mice exacerbated allergic lung inflammation^21, 22^. A similar phenotype also resulted from the expansion of filamentous fungi following antifungal treatment^23^. Antifungal-induced dysbiosis was further found to exacerbate lung allergic inflammation via intestinal resident CX3CR1^+^ phagocytic cells^24^, suggesting that gut fungi are sensed by host innate immune mechanisms that later alter homeostatic immune responses to mucosal antigens.

While informative, these studies remain difficult to interpret because fungal colonization and/or antimicrobial treatment to specific pathogen free (SPF) mice also impact the bacterial microbiome^25, 26^, which in turn could be the direct driver of the reported immune effects. Thus, the role of gut fungi as direct and/or central contributors to developmental host physiology remains coupled with the effects driven by bacteria and other ecosystem members.

The goal of this study was to determine the role of commensal fungi in host-microbiome interrelationships as they relate to microbiome assembly and host immune and physiological development. Such studies are difficult to achieve using animal models with complex or undefined microbiomes, such as SPF mice. To this aim, we have utilized a gnotobiotic approach to interrogate the capacity of commensal fungi to (i) colonize the mouse gut, (ii) alter microbiome ecology and its response to antimicrobials, and (iii) impact gut physiology and systemic immunity. We explored this in ex-germ-free (GF) mice colonized with defined consortia of either bacteria, fungi or both. For bacteria, we utilized the Oligo-MM^12^ consortium, which consists of 12 mouse-derived bacterial strains that are persistent, inheritable and eliciting an immune response in mice that is similar to a complex microbiota ^27, 28^. For fungi, we selected six yeast strains from taxa that commonly colonize the human gut and have been previously linked to atopy and asthma risk early in life^12, 13^. Our work shows that although the mouse gut can be persistently colonized with fungi, the gut is more naturally adapted to bacterial colonization. The presence of fungi elicited strong microbiome and immunological shifts that modulated subsequent susceptibility to mucosal inflammation in the distal gut and the lung. Our results provide evidence for the role of gut fungi as key players in microbiome-driven host immune development, and prompt for the incorporation of fungi in host microbiome studies.

## Results

### Fungal species colonize mouse intestinal tract less efficiently than bacteria

We first evaluated the ability of common yeast species to colonize and persist in the mouse gut. GF adult dams were colonized with defined microbial consortia of either bacteria (12 species) (B), yeast (6 species) (Y), a mixture of all the bacteria and yeasts (BY), or kept germ free (Fig. 1a, Supplementary Table 1). Dams were bred, and offspring were further colonized by spreading the dams’ abdominal area, including the nipple area, with the same inocula during the first week of life (Fig. 1a). We determined intestinal colonization and GF status by performing quantitative polymerase chain reactions (qPCR) in feces of the offspring (F1 mice) by using bacterial 16S and fungal 18S rRNA genes specific primers (Supplementary Table 1) and conventional culture. Bacteria were only detected in B and BY mice, while fungi were detected in Y and BY groups (Supplementary Fig. 1a-b). Fungal 18S RNA gene copy number was two orders of magnitude lower than bacterial DNA at 4 weeks of age (Supplementary Fig. 1a-b). At 9-10 weeks, fungal DNA copy number was ten-fold lower than bacterial DNA in the Y mice, and at this time point the 18S signal was lost in the BY group (Supplementary Fig. 1b). Given the known limitation of qPCR techniques to amplify low amplicon concentrations, we also cultured feces collected from all mice at 7 weeks of age. Fungal colonies grew from fecal cultures derived from mice in the Y and BY groups only, and fungal cell counts were ten-fold lowed in the BY group (10^6^ vs. 10^5^ CFU/ml, Supplementary Fig. 1c-d), confirming that while fungi are able to colonize the mouse gut, their fitness is reduced in this environment in comparison to bacteria, which are known to colonize ex-GF mice with at least 10^10^ CFU/ml^29, 30^.

**Fig. 1:**
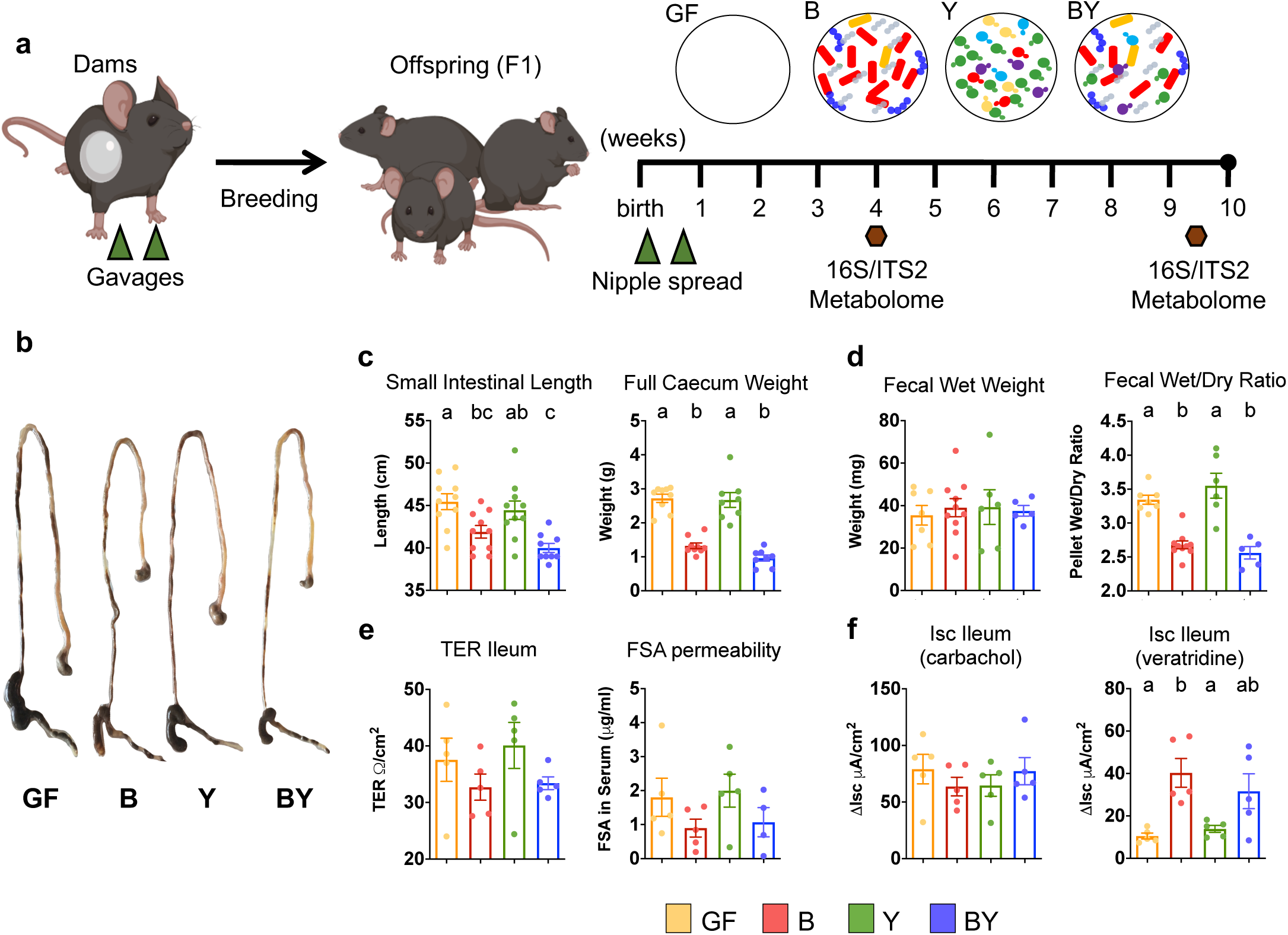
Bacterial colonization is essential to induce intestinal anatomical and functional changes of colonized mice. **(a)** Experimental layout for gnotobiotic study. Germ-free dams were orally gavaged twice with a consortium of 12 bacteria (B), 6 yeasts (Y), a combination of both (BY), or kept germ free (GF). F1 mice were further colonized during the first week of life (see methods). Fecal samples (hexagons) were obtained at 4 and 9-10 weeks of age for microbiota quantification (qPCR), taxonomic analysis (16S and ITS2 sequencing), and functional characterization (metabolome). **(b)** Representation of gross anatomic changes of dissected gastrointestinal tracts across groups (stomach to distal colon). **(c)** Small intestine length and full caecum weight of gnotobiotic dams at around 20 (17-25) weeks of age. **(d)** Water content in fecal samples measured by total fecal weight and wet/dry ratio in dams. **(e)** Gut barrier function measured by ileal transepithelial resistance (TER) and clearance of fluorescein-5(6)-sulfonic acid (FSA) over 4 hours. **(f)** Ileal short circuit current (ΔIsc) upon stimulation with epithelial stimulator carbachol or neurostimulator veratridine. (c-f) Color denotes colonization treatment (GF = yellow, B = red, Y = green, BY = royal blue); (c-d) N=6-10; (e-f) N=4-5; different letters above bars indicate statistically significant differences defined by ANOVA and Tukey posthoc tests; P<0.05.

### Fungal colonization is insufficient to trigger intestinal physiological changes

We carried out comprehensive gut physiological assessments to determine if fungal cells elicit functional changes to the organ harboring them. Anatomical measurements revealed that bacteria, but not fungi, led to a reduction in small intestinal length and full caecum weight, with Y mice exhibiting features undistinguishable from GF mice (Fig. 1b-c). Colon length and empty caecum weight remained unchanged across all groups (Supplementary Fig. 2a). Bacteria were further essential to induce changes in water absorption and intestinal permeability. Mice from groups B and BY, but not Y, displayed a reduction of fluid content in feces compared to GF (fecal wet/dry ratio; Fig. 1d). Further, examination of the clearance of fluorescein-5(6)-sulfonic acid (FSA), a measurement of intestinal barrier integrity, and ileal transepithelial electrical resistance (TER), a measure of ileal paracellular permeability, showed suggestive reductions in the mice colonized with bacteria (B and BY, Fig. 1e) but no significant differences between groups. We then evaluated the ability of the ileum to transport ions by measuring changes in short circuit current (ΔIsc) after activating enteric nerves with veratridine or directly stimulating the epithelium with carbachol (Fig. 1f). Microbial colonization did not influence ΔIsc in response to carbachol; however, GF and Y mice were almost irresponsive to veratridine (low ΔIsc), indicating that neurally-induced ion transport in the ileum relies on bacterial, but not fungal colonization. In contrast to functional changes in the ileal mucosa, no physiological differences were detected in the colon (Supplementary Fig. 2b-c).

### Exclusive fungal colonization is insufficient to promote DSS-induced colitis but exacerbated disease severity if co-colonized with bacteria

Fungal dysbiosis has been associated with IBD severity^18, 19^, although little is known about the specific contribution of common fungal species to this group of disorders. To assess if fungal colonization impacts the development of colitis, seven-week-old gnotobiotic mice colonized with the aforementioned consortia were treated with 1.5% DSS for five days (Fig. 2a). Except for one animal in the BY group, all mice survived 5 days of DSS treatment. As compared with mice colonized with bacteria (B and BY), exclusive yeast colonization resulted in decreased inflammatory features, comparable to those of DSS-untreated mice (naive) and GF mice (Fig. 2b-e), indicating that fungi alone are insufficient to trigger the immune mechanisms necessary for overt DSS-colitis. When co-colonized with bacteria, BY mice exhibited the highest colitis severity, as measured by body weight loss, spleen weight, onset of positive blood in stool, and stool lipocalin 2 (Lcn-2, a marker of neutrophil infiltration)^31^ (Fig. 2b-d). While BY mice exhibited worse disease scores, B mice displayed greater concentrations of pro-inflammatory cytokines IFN-*γ,* TNF-*α*, and IL-22 (Fig. 2e), and Y mice displayed cytokine patterns similar to GF mice (Fig. 2e). Altogether, these results suggest that while DSS-colitis relies on bacterial signals, fungal co-colonization can modulate the severity and immunophenotype of the immune response. Histopathological scoring of DSS-treated colons showed an increase in goblet cell depletion in B and BY groups compared to GF and naive animals (Supplementary Fig. 3b-c) yet no overall differences in total inflammation score for the different colonization groups (Supplementary Fig. 3a).

**Fig. 2:**
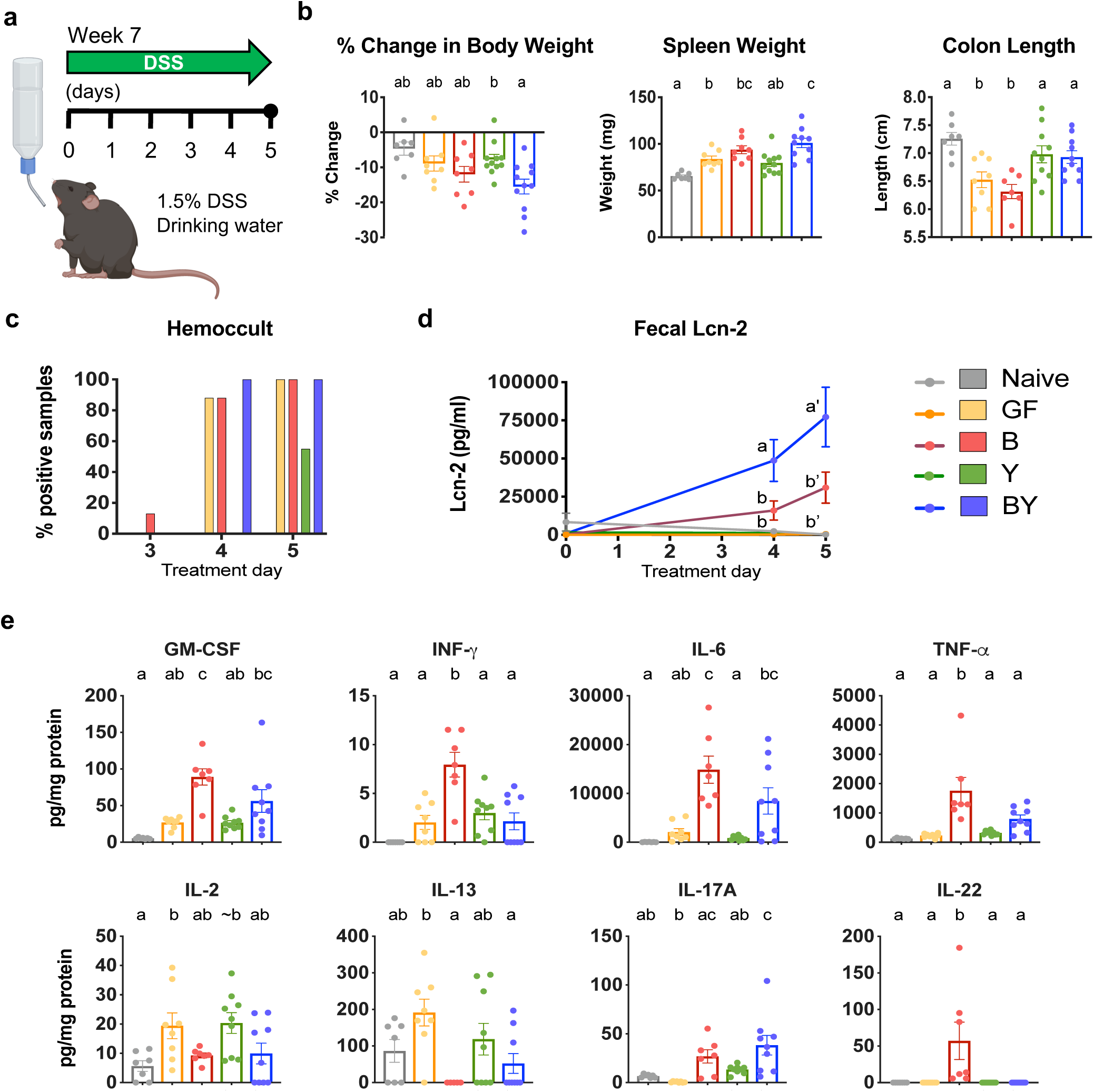
Fungal immune modulation of DSS-induced colitis. **(a)** Experimental layout for DSS study. Seven-week old gnotobiotic F1 mice were treated with 1.5% DSS for five days and colonic inflammation was assessed at end of treatment. A group of colonized animals were not treated with DSS and denoted naive (see methods). **(b)** Percentage weight change between days 1 and 5, spleen weight and colon length after 5 days of DSS treatment. Measurements of **(c)** stool occult blood and **(d)** stool Lcn-2 throughout DSS treatment. **(e)** Pro-inflammatory cytokines detected in inflamed colon lysates by electrochemiluminescence (MSD). (b-e) Color denotes colonization and DSS treatment (Naive = gray, GF = yellow, B = red, Y = green, BY = royal blue); N=7-11; different letters above bars indicate statistically significant differences defined by ANOVA and Tukey posthoc tests; P<0.05.

### Fungi induce ecological changes to the gut bacterial microbiome

We applied 16S (bacteria) and ITS2 (fungi) rRNA gene sequencing to study the impact of fungal colonization on the bacterial microbiome, and to detect interkingdom interactions. We compared bacterial and fungal diversity and composition at 4 and 9-10 weeks to evaluate microbiome shifts during early-life and adulthood, respectively. To further unravel bacterial-fungal interactions, we examined the bacterial and fungal microbiome response to community disruption, by treating two additional mouse groups with the antibiotic Augmentin^®^ (amoxicillin-clavulanate, BY+Abx) or the antifungal Fluconazole^®^ (BY+Afx) during the second week of life (see methods). We first evaluated the influence of the type of colonization (treatment) and mouse age on bacterial and fungal beta-diversity using permutational multivariate analysis of variance (PERMANOVA)^32^ on Bray-Curtis dissimilarities among samples (Supplementary Tables 2 and 3), while carefully controlling for cage effects. Treatment explained 14.43% and 39.87%, and mouse age explained 16.76% and 4.38% of the variance in the bacterial and fungal community structures, respectively (P<0.001; Supplementary Tables 2 and 3, Fig. 3a-b). This suggests both treatment and age significantly impact bacterial and fungal community structure, yet the fungal community shows a much smaller temporal shift after the early-life period compared to the bacterial community. When the microbial communities were analyzed per time point, treatment explained significant shifts in bacterial and fungal beta-diversity at 4 weeks (51.72% and 63.98% of variance, respectively, P<0.001; Supplementary Tables 2 and 3). At 9 weeks of age, the strongest driver of bacterial beta diversity was the cage effect (24.37% of variation, P=0.006; Supplementary Table 2). In comparison, the fungal community exhibited longer-lasting variations in beta-diversity due to bacterial colonization or antimicrobial disturbances (54.45% of variance, P<0.001; Supplementary Tables 3, Fig. 3a-b).

**Fig. 3:**
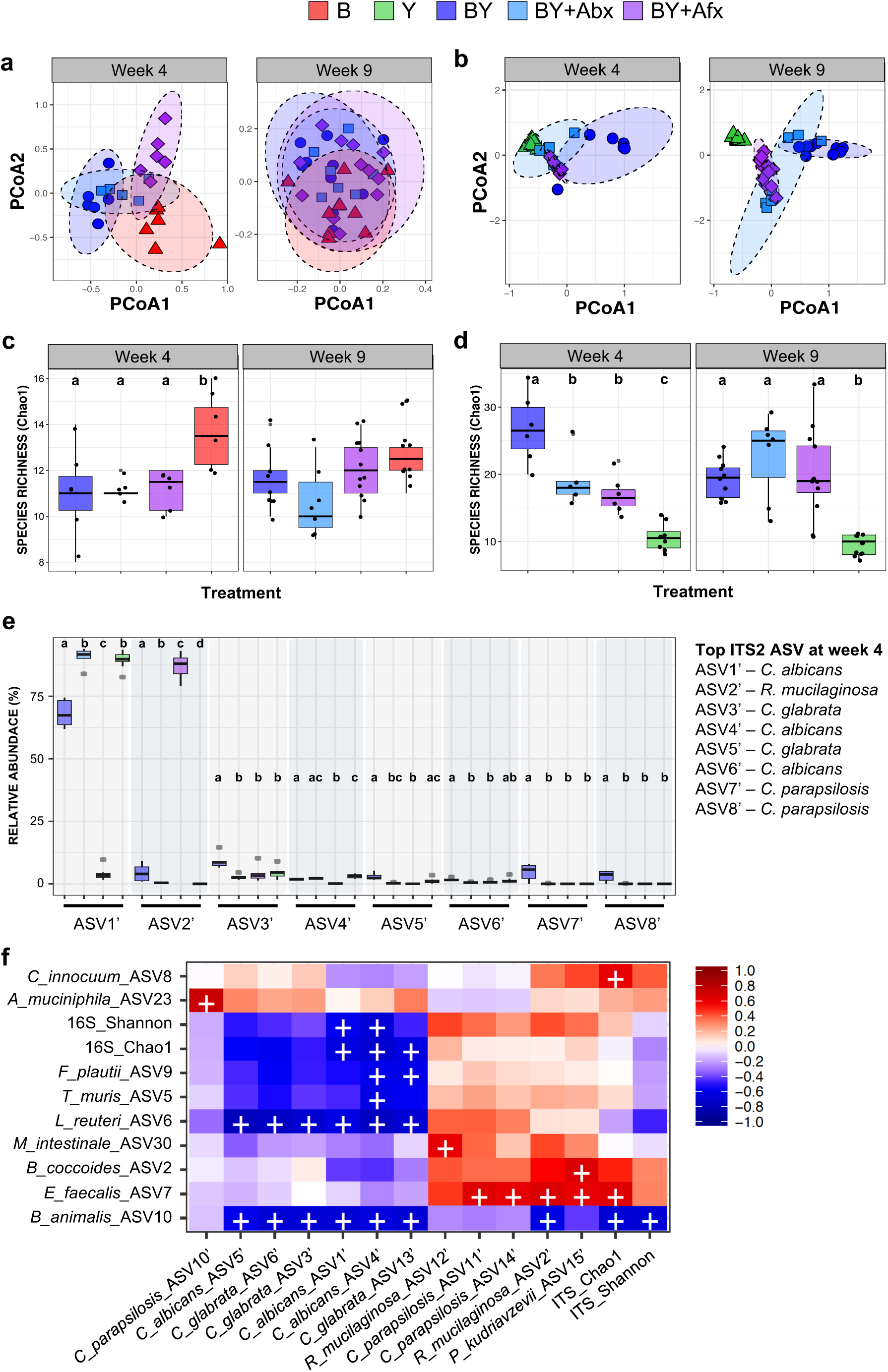
Fungal colonization and antimicrobial treatments influence gut microbiome ecology. Ecological community analyses of 16S and ITS2 rRNA sequences. PCA ordination of variation of **(a)** bacterial or **(b)** fungal beta-diversity of mice gut microbial communities based on Bray-Curtis dissimilarities among samples across treatments and experimental time points. Chao1 plots of **(c)** bacterial or **(d)** fungal species richness across groups. **(e)** Relative abundances of the 8 most abundant fungal ASVs at 4 weeks. (a-e) Color denotes colonization treatment (B = red, Y = green, BY = royal blue, BY+Abx = cyan blue, BY+Afx = purple); N^week4^=5-8; N^week9^=7-12; (c-d) different letters above bars indicate statistically significant differences defined by Kruskal-Wallis with post-hoc Dunn tests and FDR corrected; P<0.05. **(f**) Heat map of biweight correlations between top 30 bacterial (y-axis) and top 30 fungal ASVs (x-axis) in feces collected at 4 weeks of age. Color denotes positive (red) and negative (blue) correlation values. Significant correlations are denoted with a cross and defined by bicor method with FDR; P<0.05.

We also compared species richness (Chao1) and alpha-diversity (Shannon diversity index) across treatments (non-parametric Kruskal-Wallis tests followed by post-hoc Dunn tests with Benjamini-Hochberg FDR correction). Bacterial richness was reduced by fungal colonization or antimicrobial treatments at 4 weeks but not at 9 weeks of age (Fig. 3c). No differences across treatment groups were detected when evenness was also considered for either time point (Shannon index, Supplementary Fig. 4a). Interestingly, fungal alpha-diversity and richness were drastically reduced in mice harboring only fungi when compared to BY mice at 4 weeks of age, reaching lower diversity than in mice that were treated with antimicrobials (Fig 3d, Supplementary Fig 4b), indicating that gut fungal diversity benefits from bacteria during early-life colonization. Fungal alpha-diversity increased by week 9 in all groups except in the antifungal challenged community (Supplementary Fig. 4b), whereas fungal richness remained reduced for the Y only mice (Fig. 3d). Altogether, these results show that bacterial and fungal population reciprocally change each other, with stronger and longer lasting effects of bacteria on the fungal community. These inter-kingdom effects were comparable to the effects of antimicrobials.

To identify the microbial taxa driving these community shifts we determined the differential relative abundance of the top amplicon sequencing variants (ASV) across treatment groups (non-parametric Kruskal-Wallis tests followed by post-hoc Dunn tests with Benjamini-Hochberg FDR correction). For the bacterial microbiome, most changes attributed to fungal colonization impacted less predominant taxa, including *Enterococcus faecalis, Clostridium innocuum, Bacteroides caecamuris,* and *Lactobacillus reuteri*. Fungal colonization distinctly antagonized the growth of *L. reuteri*, which was drastically decreased in BY mice compared to B or B+Afx mice (Supplementary Fig. 4c).

We detected stronger compositional changes for all eight of the most abundant fungal ASVs (Fig. 3e). The top two ASVs, *C. albicans* and *Rhodotorula mucilaginosa* represented over 90% of the fungal communities and exhibited the largest shifts across treatments. *C. albicans* dominated the community in all groups, except in mice treated with fluconazole, in which *R. mucilaginosa* was the dominant taxa (Fig. 3e), suggesting interspecies competition or resistance to fluconazole *in vivo* (all fungal strains were susceptible to fluconazole *in vitro*). Co-colonization with bacteria resulted in increased abundance of *Candida* species, *C. glabrata* and *C. parapilopsis*, whereas *C. albicans* dominated in the absence of bacteria or in antibiotic-treated mice (Fig. 3e).

Correlative analysis of the relative abundances of the 30 most abundant ASVs of the 16S and ITS2 datasets provided further insight into inter-domain ecological interactions at play in these defined communities (bicor method, FDR corrected). We identified strong positive correlations between *A. muciniphila* and *C. parapilopsis*, as well as between *E. faecalis* and *C. parapilopsis*, *R. mucilaginosa* and *P. kudriavzevii* (Fig. 3f). Negative correlations were detected between two bacterial species *L. reuteri* and *B. animalis*, and three species of *Candida* and *R. mucilaginosa* (Fig. 3f). Interestingly the abundance of certain species impacted alpha-diversity measures of the opposite domain, such as between *E. faecalis* and *B. animalis* vs. fungal Chao1 and Shannon indexes, as well as *C. albicans* and *C. glabrata* vs. bacterial alpha diversity and richness (Fig. 3f). Altogether, our findings convincingly show that fitness of some of the most common yeast species in the mouse gut benefit from the presence of bacteria, while the growth of some bacterial strains, such as *L. reuteri,* is antagonized by fungal colonization.

### Fungal colonization has limited impact on fecal metabolome

Microbial metabolites produced in the gut are predicted to modulate immune development, though very little is known about the contribution of fungal species. We explored the functional changes attributed to fungal colonization by determining the fecal metabolomes for each of our gnotobiotic groups by liquid chromatography coupled to mass spectrometry (LC-MS). Principal component analysis (PCA) performed on the relative amounts of 150 metabolites, which were obtained from feces collected at week 4 and 9, demonstrated that the metabolome profile of Y mice was similar to GF, which were both very different from the groups colonized with bacteria, B, BY, BY+Abx, and BY+Afx (Fig. 4a). Statistical analysis revealed 99 metabolites with significantly different amounts between the groups (Supplementary Table 4; ANOVA+FDR; P<0.05). Heat-map of the top 25 most differentially detected metabolites shows clustering of Y and GF samples separately from bacteria-colonized groups (Fig. 4b). Metabolites detected at lower levels in the GF and Y groups were associated with the metabolism of butanoate, ubiquinone, pantothenate, coenzyme A, and mostly aromatic amino acids (Supplementary Table 4). The presence of bacteria reduced levels of several oligo and monosaccharides and a large number of amino acids, indicating bacterial consumption of these compounds (Supplementary Table 4). These results thus suggest that fungal metabolites do not contribute significantly to overall fecal metabolome of mice colonized with both bacteria and fungi.

**Fig. 4:**
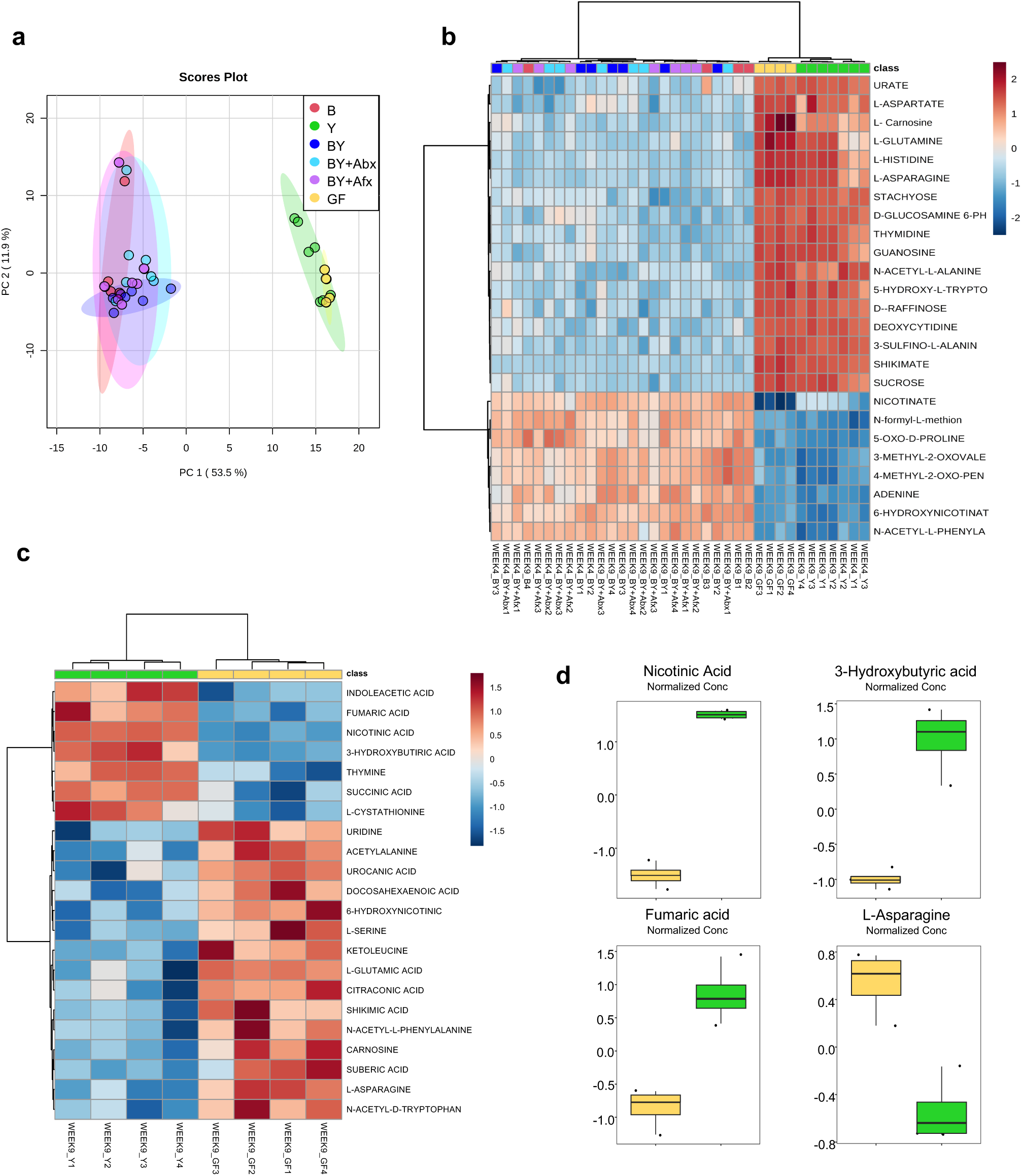
Bacteria colonization is the main influencer of changes in the fecal metabolome. **(a)** Principal component analysis score plot of 150 metabolites detected in fecal samples of gnotobiotic mice at 4 and 9 weeks of age. **(b)** Heat map of top 25 differentially expressed metabolites detected in 4- and 9-week fecal samples. **(c)** Heat-map of 22 metabolites differentially expressed between GF and Y groups at 9 weeks. **(d)** Strongest metabolic differences detected between GF and Y groups at 9 weeks. (a-d) Color denotes colonization treatment (GF = yellow, B = red, Y = green, BY = royal blue, BY+Abx = cyan blue, BY+Afx = purple); N^week4^=3; N^week9^=4. (a-b) Statistically significant differences between groups determined by ANOVA with Fisher’s post-hoc and FDR; P<0.05. (c-d) Statistically significant differences defined by two-tailed t-test and FDR corrected; P<0.05.

Although the metabolome profiles of the Y and GF groups appeared very similar, we identified 22 metabolites with significant different levels within Y and GF samples at 9 weeks (Fig. 4c, Supplementary Table 5; T-test+FDR; P<0.05). Y group showed increased levels of metabolites produced in citric acid cycle and butanoate metabolism, while GF mice had significantly elevated levels of metabolites associated with metabolism of histidine, asparagine, serine and nitrogen (Supplementary Table 5). The most significant changes originating from fungal colonization were detected for nicotinic acid, 3-hydroxybutyric acid, fumaric acid and L-asparagine (Fig. 4d, Supplementary Table 5). This confirmed that the colonizing fungi are metabolically active in the murine intestinal tract.

We also looked at metabolome profiles of the small intestine content for B, BY, Y and GF mice at week 4 (Supplementary Fig. 5). As opposed to the fecal metabolome data, neither PCA of 124 quantified metabolites (Supplementary Fig. 5a) nor hierarchical clustering analysis of the relative abundances for the top 25 detected metabolites (Supplementary Fig. 5b) yielded specific grouping of the mice. This observation agrees with the fact that highest bacterial load is located in the large intestine, with small intestine harboring generally lower bacterial amounts^33^. In conclusion, although fungal colonization promotes strong alterations in the bacterial microbial community, its impact on the fecal metabolome profile of colonized mice is limited.

### Fungal colonization alters early-life systemic and gut mucosal immunity

To examine the immune consequences of fungal colonization, we characterized the immune response at 4-weeks of age by using flow cytometry of extracellular and intracellular markers in unstimulated splenocytes (flow cytometry gating strategy outlined in Supplementary Fig. 6), immunoglobulin (Ig) quantification in serum, and multi-array cytokine determination in jejunum and colon lysates (see methods). Fungal colonization promoted broad systemic immunological changes to immune cell populations and their secretion of cytokines in the spleen (Fig. 5a-b, Supplementary Fig. 7a-b). BY mice exhibited shifts in CD19^+^ B cells, CD3^+^ T cells, FoxP3^+^ regulatory T cells (Tregs), and Lin^−^ (CD19^−^CD3^−^) myeloid cells, compared to B and GF groups (Fig. 5a). Antibiotic treatment resulted in similar changes in B and T cell populations, compared to B and GF mice (Fig. 5a). Within the Lin^−^ population we also detected alterations in macrophage subpopulations. BY-colonization increased the proportion of F4/80^+^MHC-II^+^CD11c^−^ macrophages (Fig. 5a). Changes in Tregs and macrophage subpopulations are likely attributed to fungi as Y and BY mice exhibited the same patterns, compared to GF and B mice (Fig. 5a). Antimicrobial treatments to BY mice resulted in changes in most of the splenocyte populations, compared to untreated BY mice (Fig. 5a and Supplementary Fig. 7a). Systemic immune changes were also evident from cytokine secretion by splenic cells and serum antibodies. Percentages of splenocytes producing IL-4, IL-6, IL-10 and IL-12 cytokines in BY groups were significantly increased from GF or B (Fig. 5b), suggesting a synergistic immune effect when a host is colonized with fungi and bacteria. Increased IL-4 and IL-6 secretion was mainly driven by CD4^+^ T and Lin^−^CD11c^+^ cells, while Lin^−^CD11b^+^ cells displayed increased expression of IL-6 and IL-12 (Supplementary Fig. 7b). Exclusive fungal colonization resulted in decreased serum IgG3, as well as increased IgG1 (Fig. 5c), and a non-significant increase in IgE compared to B group (Supplementary Fig. 7c). As well, BY and BY+Abx mice displayed increased levels of IgA and IgG3 (Fig. 5c). Importantly, these striking systemic immune changes were not a result of systemic fungal infection as fungal cultures of the kidneys (a classic marker of fungemia)^34^ were negative (data not shown).

**Fig. 5:**
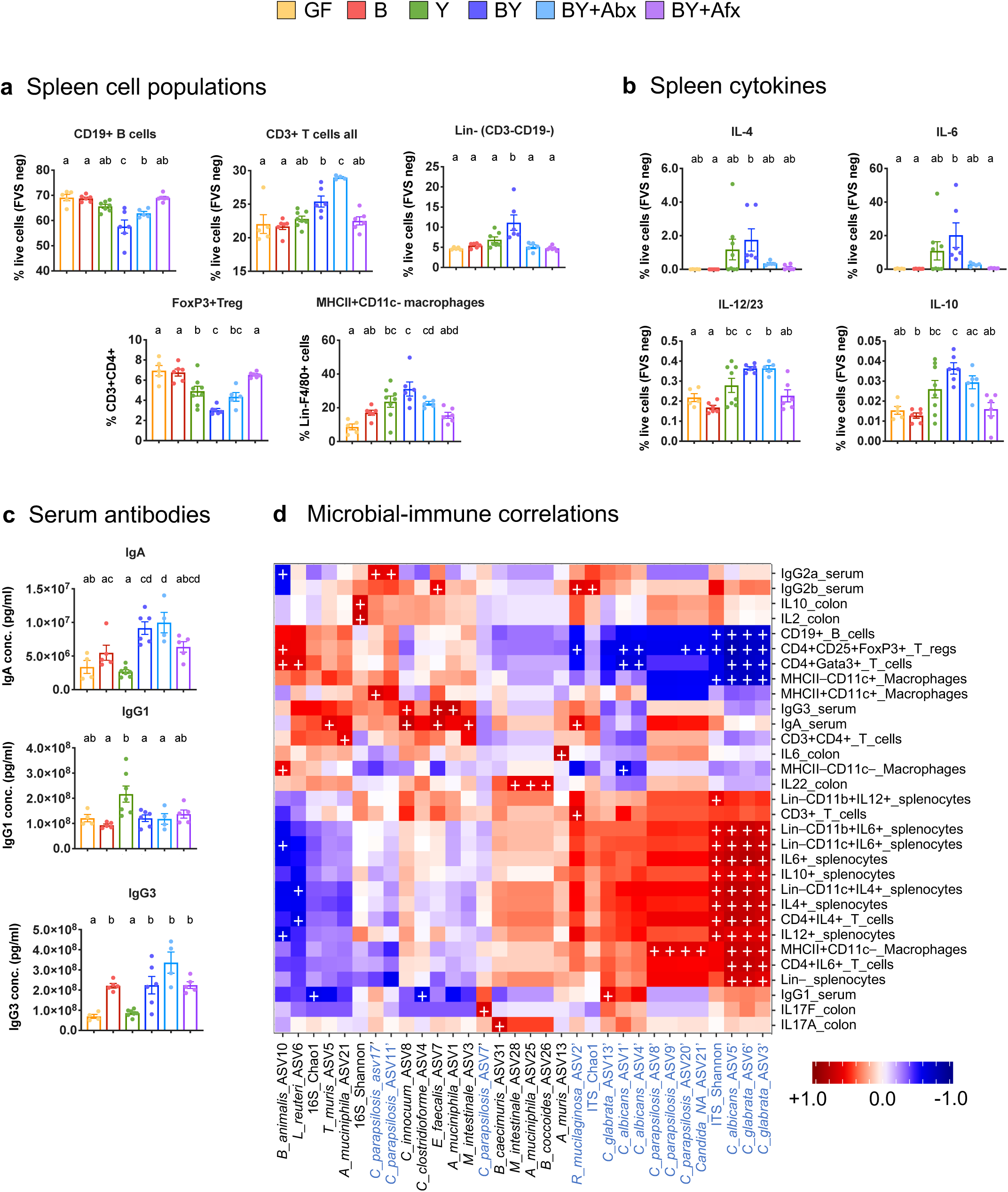
Fungal colonization alters early-life systemic immunity in mice. Percentage of **(a)** unstimulated splenic cell populations, and **(b)** cytokine-producing unstimulated splenocytes from 4-week-old gnotobiotic mice. Cells were stained with intra- and extracellular marker-specific antibodies and quantified by flow cytometry. **(c)** Serum antibody concentrations detected by electrochemiluminescence (MSD). (a-c) Color denotes colonization treatment (GF = yellow, B = red, Y = green, BY = royal blue, BY+Abx = cyan blue, BY+Afx = purple); N= 5-8; different letters above bars indicate statistically significant differences defined by ANOVA and Tukey posthoc tests; P<0.05. **(d)** Heat map of correlations between 16S (black) and ITS (blue) ASVs (x-axis) and reported immune features (y-axis) at week 4. Color denotes positive (red) and negative (blue) correlation values. Significant correlations are denoted with a cross and defined by bicor method with FDR; P<0.05.

Microbial colonization also drove mucosal immune changes in the jejunum and colon (Supplementary Fig. 8). Cytokine levels in jejunum lysates of GF mice consistently showed high levels of pro-inflammatory cytokines, IL-6, IL17A, IL-21, INF-g, TNF-alpha, and GM-CSF (Supplementary Fig. 8a). Microbial colonization decreased these cytokine levels, but only with bacterial colonization (B and BY mice) whereas exclusive colonization with fungi was insufficient to induce this effect, except for IL-21 (Supplementary Fig. 8a). Microbial colonization also affected cytokine level in the colon but fewer changes were attributed to fungal colonization (Supplementary Fig. 8b).

### Systemic immune alterations are strongly correlated to fungal colonization

To further characterize the specific microbial drivers of the aforementioned immunological changes, we performed correlated the relative abundance of bacterial and fungal ASVs with the reported immune features (bicor method, FDR corrected). Remarkably, most of the significant correlations observed were associated to fungal colonizers (Fig. 5d). *C. albicans* and *C. glabrata* species promoted the most striking systemic changes observed, including the increased proportions of IL-4, IL-6, IL-10 and IL-12-producing splenocytes, as well as proportions of B cells, Tregs, Gata3-producing T cells, and MHC-II^−^CD11c^+^ macrophages (Fig. 5d). *C. parapsilosis* was also strongly correlated to an increase in MHC-II^+^CD11c^−^ and MHC-II^+^CD11c^+^ macrophages, reduction of Tregs, and increase in serological levels of IgG2a (Fig. 5d). Fungal community alpha-diversity and richness also significantly impacted systemic immunity. Fungal alpha-diversity (Shannon) was positively correlated to increased proportions of pro-inflammatory cytokine producing cells, and reduction of B cells, Tregs and MHC-II^−^CD11c^+^ macrophages (Fig. 5d). Fungal richness (Chao1) was associated to increased T cell and reduced Treg proportions, and increased serum IgA and IgG2b (Fig. 5d).

We detected more discrete correlations between some of the bacterial colonizers and immune markers. *B. longum* subsp. *animalis* was correlated with reduced secretion of IL-6 and IL-12 pro-inflammatory cytokines, and increased proportions of MHC-II^−^CD11c^−^ macrophages, Tregs and Gata3-producing CD4^+^ T cells (Fig. 5d). Similarly, *L. reuteri* was correlated with reduced secretion of IL-4 cytokine while increasing Gata3^+^ CD4^+^ T cells (Fig. 5d). The most abundant colonizer *A. muciniphila* was associated with an increased secretion of IL-2, IL-10 and IL-22 in colon lysates, and increased IgG3 levels in serum (Fig. 5d). Bacterial community Shannon and Chao1 indexes were also associated to increased proportion of CD4^+^ T splenocytes and serum IgG1, respectively (Fig. 5d).

We also applied sparse generalized canonical correlation analysis (SGCCA) to further determine the relevance of the top 15 colonizers from the 16S and ITS datasets on the measured immunological features. Relevance network plots identified *C. albicans* and *R. mucilaginosa* as the two fungal taxa with most relevant correlations with systemic immune cell populations (Supplementary Fig. 9). This method confirmed many of the above-mentioned correlations (Fig. 5d), as well as a negative correlation between *C. albicans* and jejunum IL-2 levels (Supplementary Fig. 9). SGCCA also detected bacterial species, *C. innocuum, C. clostridioforme, M. intestinale, B. coccoides* and *A. muciniphila* as relevant for splenocyte populations and their cytokine responses (Supplementary Fig. 9). Altogether, these results highlight that fungal colonization is strongly correlated with the observed systemic immunological features.

### Gut fungal colonization modulates OVA-induced airway inflammation and infiltrate phenotype

Fungal alterations are associated with subsequent asthma in humans^12^, yet causality remains to be determined. We explored the impact of fungal colonization on host susceptibility to airway inflammation by challenging mice with ovalbumin (OVA; Fig. 6a, see methods). We observed that GF mice were highly susceptible to OVA-airway inflammation, demonstrating increased bronchoalveolar lavage (BAL) cellularity and eosinophilia, relative to naive (non-OVA challenged) animals (Fig. 6b), as previously described^35^. Compared to naive mice, exclusive fungal colonization and antimicrobial treatments in BY+Abx and BY+Afx groups also resulted in elevated measures of lung inflammation (Fig. 6b).

**Fig. 6:**
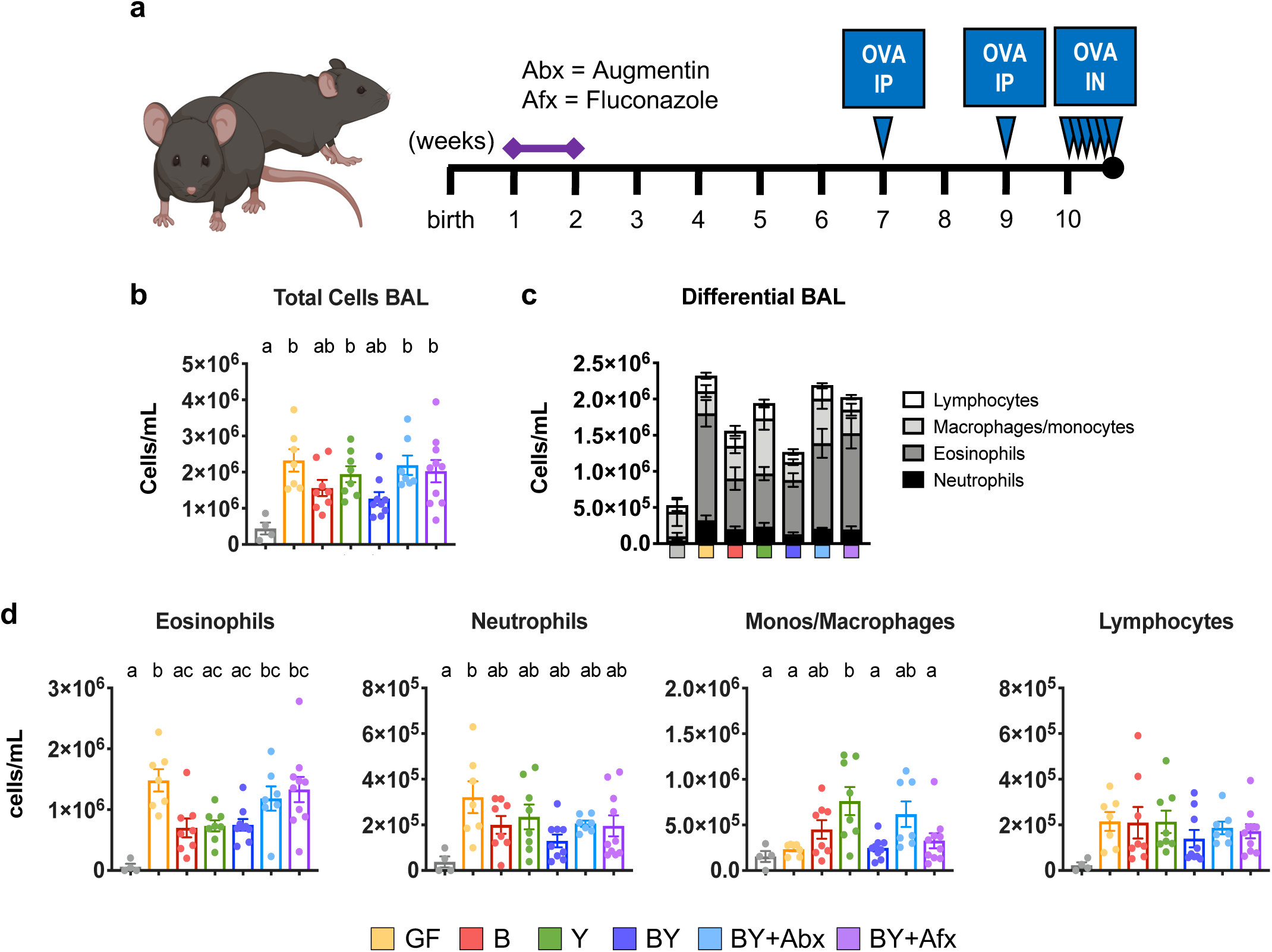
Fungal colonization modulates OVA-induced airway inflammation phenotype. **(a)** Experimental layout for OVA-induced airway inflammation. Two groups of BY animals were additionally treated with antimicrobials Augmentin (Abx) or Fluconazole (Afx) during the second week of life. F1 mice were systemically sensitized with two intraperitoneal (IP) injections of OVA+Alum at weeks 7 and 9. Afterwards, F1 mice were challenged intranasally (IN) with OVA, for 5 consecutive days at week 10. A group of mice were kept unchallenged and denoted naive (see methods). **(b)** Total cellular count in BAL fluid following OVA challenge. **(c)** Compiled differential counts of cellular infiltrate in gnotobiotic groups. **(d)** Differential leukocyte counts in challenged mice. (b-d) Color denotes colonization and OVA challenge (Naive = gray, GF = yellow, B = red, Y = green, BY = royal blue, BY+Abx = cyan blue, BY+Afx = purple); N=5-10; different letters above bars indicate statistically significant differences defined by ANOVA and Tukey posthoc tests; P<0.05.

Differential cellular counts of the inflammatory infiltrate highlighted the distinctive phenotype and tone of the inflammatory responses. Increased eosinophilia was observed in BY+Abx and BY+Afx groups, while the increased cellular infiltrate in the Y group was marked by increased counts of monocytes/macrophages (Fig. 6c-d), suggesting that fungal colonization can modulate the immune phenotype of airway inflammation. Overall, the presence of bacteria (B and BY) but not fungi reduced acute airway inflammation in this model, suggesting that homeostatic control of lung allergic responses relies on bacterial signals.

## Discussion

Using a gnotobiotic approach, we tested the direct and indirect causal effects of fungal colonization on intestinal physiology and systemic immune development, as well as their ecological role in the microbial community. To our knowledge, this is the first time the role of fungi has been investigated in the absence of bacteria, enabling the study of their direct contribution to host development. Our work provides a mechanistic analysis on the role of fungi, showing that while the mouse gut is more physiologically equipped to harbor bacteria, fungi strongly impact microbiome dynamics (Fig. 3, Supplementary Fig. 4), and promote robust local and systemic immunological changes (Fig. 5, Supplementary Fig. 7, Supplementary Fig. 8, Supplementary Fig. 9) that influence the immune phenotype of intestinal and lung inflammatory responses (Fig. 2, Fig. 6).

Previous work has suggested that fungi are unable to colonize the intestinal tract of healthy humans, proposing that fungal detection in fecal samples originates from fungal transiting through the gastrointestinal tract from food sources or the oral cavity^36^. In contrast, our mouse model consistently showed that, even with periodic cage bedding changes and sterile food sources, we were able to culture fungi from gnotobiotic fecal samples several weeks after initial inoculation (Supplementary Fig. 1c-d). These results indicate that gut fungi are most certainly gut dwellers, albeit in far lower concentrations than gut bacteria. Successful fungal colonization in the absence of bacteria further indicates that the latter are not required for fungi to engraft in the mouse gut. Fungi grew in higher concentration in the absence of bacteria, in line with previous findings showing interkingdom competition or antagonism between bacteria and fungi in other ecosystems, such as the rhizosphere and soil^37^.

While not reflective of natural conditions, exclusive fungal colonization is a valuable experimental benchmark to differentiate the effects of bacteria and fungi, and potential synergism or antagonism in these effects. This condition was essential to determine that although fungi colonized the mouse gut, their presence was insufficient to induce any physiological or morphological changes in the ex-GF mice. Intestinal anatomy and physiology in mice exclusively colonized with fungi were essentially identical to the GF mice (Fig. 1, Supplementary Fig. 2), suggesting that the reported adaptations of the gut tissue to microbial colonization ^33^ may not extend to fungal organisms.

Similarities in cytokine response between GF and Y groups were also observed following five days of treatment with DSS, in which absence of microbes or exclusive fungal colonization resulted in reduced inflammation. This supports previous work showing that bacterial colonization is essential for the development of DSS-colitis^38^, and expands current knowledge by determining fungal colonization as insufficient to induce colitis. In line with previous studies, our model showed higher expression of IL-17A in intestinal tissues in bacteria-colonized groups following DSS treatment (Fig. 2e). IL-17A is a potent enhancer of DSS-induced colitis phenotype, with elevated IL-17A secreted by Th17 cells promoting secretion of IL-12 and IL-23, and triggering a conversion of the response from Th17 to Th1^39, 40^. Our model showed that bacteria colonization induced elevated detection of Th1 pro-inflammatory cytokines INF-γ, IL-6 and TNF-α, which have been previously suggested to create an inflammatory loop and worsen colitis in mice^41^. In addition, our work convincingly showed that IL-22 secretion only occurred in exclusive colonization with bacteria, and that presence of fungi prevented this (Fig. 2e), which may help explain the changes in inflammatory response attributed to fungal colonization. IL-22 plays dual roles in intestinal inflammation. For instance, T cell production of IL-17 and IFN-γ is associated with a deleterious function of IL-22^42^, yet colonic delivery of IL-22 gene via a secretory vector increased goblet cell expression and ameliorated DSS-induced colitis^43^. In this context, the prevention of IL-22 secretion in mouse groups colonized with fungi (Y and BY) may be a contributing factor to the elevated inflammatory response in in co-colonized mice (BY). Further research testing the modulatory capacity of IL-22 in gnotobiotic conditions will be necessary to confirm this.

Several studies have reported that fungal overgrowth exacerbates DSS-colitis in SPF mice^15, 44, 45^. However, in light of our observations that fungal signals alone are unable to induce the level of colonic inflammation commonly reported in DSS-treated animals (Fig. 2), it is more likely that interactions bacteria and fungi leads to exacerbated colitis. A recent study in mice provides convincing evidence that CARD9 signaling via Dectin-2 receptor activation in *Malassezia*-treated animals worsens colitis^19^. In their work, Limon *et al.* demonstrated that CARD9 activation induced a pro-inflammatory cascade towards innate immune cell activation and exacerbation of intestinal inflammation^19^. Further research is needed to confirm if this activation is also dependent on the concomitant presence of bacterial signals, as our results suggest, or if it is driven by specific fungal species, as our fungal consortium did not include *Malassezia sp*.

Our data also bring to light that the gut environment is conducive to very strong interkingdom bidirectional interactions. The presence of fungi impacted bacterial community structure, although bacteria had a stronger and longer lasting effect on the fungal community (Fig. 3, Supplementary Fig. 4). Co-colonization with bacteria strongly altered the fungal community’s beta-diversity, and significantly increased alpha-diversity and richness (Fig. 3, Supplementary Fig. 4). Interestingly, the impact of fungal colonization on bacterial beta-diversity and richness was stronger at 4 weeks of age (Fig. 3), supporting the concept of a window of opportunity early in life, during which the microbiome is more susceptible to alterations^7^.

Microbial colonization early in life is critical for immune education^46, 47^, and our work provides precise evidence that fungi, in the absence or presence of bacteria, stimulates systemic immune responses during early life (Fig. 5, Supplementary Fig. 7). Most systemic immune changes resulted from the synergistic effects of both bacterial and fungal colonization (Fig. 5a-b, Supplementary Fig. 7a-b), including shifts in all major immune cell subtypes (B, T, Tregs, macrophages, Lin^−^ cells), as well as increased cytokines (IL-4, IL-6, IL-12 and IL-10), suggesting a systemic stimulatory effect in co-colonized animals. The importance of the early-life bacterial and fungal microbiome was further supported by studies in mice treated with antimicrobials, with antibiotic- and antifungal-treated mice displaying similar trends to the Y-only and B-only groups, respectively (Fig. 5a-b, Supplementary Fig. 7a-b). Of note, the substantial immune changes were not due to fungal infection in any of the groups, as corroborated by negative fungal cultures in kidney, a feature of fungemia. Our results strongly suggest that commensal colonization with fungi has systemic stimulatory effects in the host. This is in line with a recent study in human T memory cells specific to *C. albicans*, which displayed cross-reactivity towards *Aspergillus fumigatus* in the airway^48^, suggesting that gut fungi colonization is important for immune education and immunological memory towards fungal infections at other mucosal sites.

Which fungal-driven mechanisms causing the abovementioned immune changes remain unknown. Our untargeted metabolomic analysis revealed that the gut-colonizing fungi are metabolically active (Fig. 4c-d). However, because we did not detect any metabolites that differed between the BY from the B-only groups we speculate that these effects are not driven by the uptake of fungal-derived metabolites by immune cells. An alternative, more plausible mechanism may involve direct interactions between fungal-associated molecular patterns with immune cells. Further mechanistic work evaluating the effect of specific structural and/or secreted fungal molecular patterns has great potential to reveal microbial patterns that may act as specific triggers of these immune responses.

Early-life fungal dysbiosis during the first months of human life has been associated with increased risk of atopy and asthma risk in children^12, 13^, and patients with severe asthma often benefit from antifungal treatments as a therapeutic approach^49^. Although these studies showed striking associations between microbial fungal community composition and airway inflammation, causality has not been directly assessed in a sufficiently controlled manner. Our results revealed that colonization with bacteria (B and BY) but not exclusive fungal colonization reduced OVA-induced airway inflammation in mice, and that fungi promote infiltration of macrophages/monocytes in the bronchoalveolar space (Fig. 6), coinciding with our findings on fungi-driven early-life changes in splenic cell populations (Fig. 5). A possible mechanism may involve prostaglandin E^2^ (PGE2), a strong regulator of macrophage polarization produced and secreted by *Candida* species. This is supported by previous work from Kim *et al.*^22^, showing that antibiotic-induced fungal overgrowth in the gut led to increased M2-macrophage infiltration and PGE2 plasma concentrations.

These findings show that bacterial signals early in life are critical to prevent increased susceptibility to airway allergic inflammation, and that fungal commensals can act as modulators of the underlying immune profile of airway inflammatory responses. Our work provides a clear portrayal of how the initial composition of the gut microbiome, as well as perturbations to it, distinctly dictate the strength and cell types involved in the immune response towards a mucosal antigen.

Overall, our unique experimental approach enabled the demonstration that fungal colonization, although insufficient to promote intestinal physiological changes in the host, significantly impacts early-life host immune development and susceptibility to inflammatory-disease in the distal gut and the lungs. This work causally implicates fungi as microorganisms that can skew the immature immune system and increase susceptibility to immune-mediated disorders later in life. Further efforts are required to fully understand the specific implications of fungal dysbiosis and, more importantly, how to prevent its manifestations.

## Methods

### Colonization of Gnotobiotic Mice

Ten to fifteen-week old GF C57Bl/6J mice were obtained from the gnotobiotic mouse facility of the International Microbiome Centre (IMC) at the University of Calgary. Female mice were orally gavaged twice, 3 days apart, with 100 μl of a consortium of microorganisms, or kept germ-free. Colonization consisted of consortia of (i) twelve mouse-derived bacteria, “B”^27^, (ii) six yeast species previously linked to atopy and asthma risk in infants, “Y” ^12, 13^, or (iii) a combination of all 18 bacteria and yeast species, “B+Y”. Inocula were prepared under anaerobic conditions by mixing 100 μl of 2-day old microbial cultures of each species. Bacteria were grown in fastidious anaerobe broth (LabM, Heywood, Lancashire, UK) and yeast were grown in yeast-mold broth (YM, BD, Sparks, MD, USA). After the second gavage, mice were paired for mating on a 2:1 female:male ratio per cage. To promote microbial colonization with the desired consortia in F1 mice, the corresponding inocula were spread on the dams abdominal and nipple regions on days 3 and 5 after birth, as previously described^20^. Two groups of mice colonized with both bacteria and yeast were also treated with the antibiotic Augmentin (0.2 mg/ml, Sigma, Oakville, ON, Canada) or antifungal Fluconazole (0.5 mg/ml, Sigma) in sterile drinking water from day 7 to 14 after birth. Treatment solutions were prepared by dissolving the antimicrobials in distilled water, followed by filter sterilization. All experiments were conducted under protocols approved by the Health Sciences Animal Care Committee, following the guidelines of the Canadian Council on Animal Care.

### Fungal Culture in Selective Medium

Fungal colonization was assessed by culture in selective medium. Mice fecal pellets (∼20 mg) were dissolved in 0.9% sterile saline, serially diluted, and plated in YM agar plates supplemented with gentamycin (10 μg/ml, Sigma) and chloramphenicol (200 μg/ml, Sigma). Plates were incubated at 25°C and checked for yeast-like colonies for up to 10 days. Fungal concentrations were determined by plate count method.

### Fungal and Bacterial Quantification (qPCR Assays)

Fungal and bacterial DNA concentrations in feces were determined by quantitative amplification of the fungal 18S and bacterial 16S rRNA genes, as previously described^12, 50, 51^. Gut microbial DNA was obtained from fecal samples using the DNeasy PowerSoil Pro kit (Qiagen, Hilden, Germany) according to the manufacturer’s instructions, eluted in nuclease free water, and stored at −80°C. All qPCR reactions were carried using iQ SYBR Green Supermix (BioRad Laboratories, Hercules, CA, USA). For 16S qPCR protocol, 10-μl reactions consisted of 5 μl master mix, 0.5 μl of each primer (u16Sfr and u16Srv, see Supplementary Table 1) at 10 μM, 2 μl nuclease free water, and 2 μl of 1 ng/μl dilution of extracted template DNA. The 16S qPCR thermocycler program consisted of an initial 5 min step at 94°C, followed by 40 cycles of 94°C for 15 s, 60°C for 30 s and 72°C for 30 s, and a final melt curve consisting of a single cycle of 95°C for 15 s, 60°C for 1 min, 95°C for 15 s, and 60°C for 15 s. For fungal 18S qPCR protocol, 20-μl reactions consisted of 10 μl master mix, 2.5 μl of each primer (FR1 and FF390, see Supplementary Table 1) at 10 μM, 3 μl nuclease free water, and 2 μl of 1 ng/μl dilution of template DNA. The fungal 18S qPCR thermocycler program consisted of an initial 10 min step at 95°C, followed by 40 cycles of 95°C for 15 s, 50°C for 30 s and 70°C for 60 s, and a final melt curve consisting of a single cycle of 95°C for 15 s, 60°C for 1 min, 95°C for 15 s, and 60°C for 15 s. All samples were run in duplicate and concentration calculated based on standard curves generated using bacterial genomic DNA (extracted with DNeasy PowerSoil Pro kit, Qiagen) or fungal genomic DNA (extracted with YeaStar Genomic DNA Kit; Zymo Research, Irvine, CA, USA). qPCR runs were performed on the StepOne Plus Real-Time PCR System (Applied Biosystems, Foster City, CA, USA) in the Snyder Resource Laboratories, University of Calgary.

### Fecal Water Content

Fecal pellets were collected and immediately weighed (5-10 mice/treatment group). Pellets were dried overnight at 50°C, reweighed to determine wet:dry weight ratios.

### Gastrointestinal Barrier Assessment and Small Intestinal Transit

Intestinal permeability was measured using previously described methods with some modifications^52^. Briefly, mice (4-5/treatment group) were gavaged orally with 100 μl of 50 mg/ml fluorescein-5(6)-sulfonic acid, trisodium salt (FSA, 478 daltons; Setareh Biotech, LLC, Eugene, OR, USA). Small intestinal transit was assessed as previously described in detail^53^. Briefly, 3hr 45min following FSA gavage mice were gavaged with 200 μl Evans Blue (5% suspended in 5% Gum Arabic; Sigma). Four hours after FSA gavage and 15 min after Evans Blue gavage animals were anesthetized with isofluorane and blood was drawn by cardiac puncture for serum FSA content. Blood was allowed to clot at room temperature, spun at 2000 g for 10 min and the supernatant (serum) read on a spectrophotometer at 485/535 nm. Serum FSA (μg/ml) was determined from a standard curve. To assess small intestinal transit animals were euthanized by cervical dislocation and the entire gastrointestinal tract from the stomach to the anus was immediately removed. The distance travelled by the colored marker was measured and expressed as a percentage of the total small intestinal length from the pylorus to the cecum. As described below, cecal weight, small intestinal and colon length were determined, and tissue from the colon was used to conduct *ex vivo* measurement of ion secretion and transepithelial resistance.

### Colonic Propulsion, Cecal Weight, and Small Intestinal and Colon Length

Distal colonic propulsion was measured as previously described^54^, with slight modifications. Animals (4-5/treatment group) were allowed to acclimate for 30 min after being transferred to the lab. Mice were lightly anesthetized with isofluorane before a plastic bead (2.5 mm diameter) was inserted 3 cm into the distal colon using a silicone pusher. The time to expulsion of the bead was determined in seconds. Following bead expulsion mice were anesthetized with isofluorane and killed by cervical dislocation and the entire gastrointestinal tract from the stomach to the anus was immediately removed for measurement of cecal weight, small intestinal and colon length. Tissue from the ileum was used to conduct *ex vivo* measurement of ion secretion and transepithelial resistance as described below. For each mouse killed (8-10/treatment group) the full cecum was weighed and then the contents removed, and the empty cecum reweighed. Both the length of the ileum, from the pylorus to the cecum, and the colon length were measured.

### Measurement of Ion Secretion and Transepithelial Resistance

Full-thickness segments of terminal ileum, proximal colon and distal colon were opened along the mesenteric border, cleaned of luminal contents, and mounted in Ussing Chambers (Physiologic Instruments, San Diego, CA, USA) with an exposed area of 0.3 cm^2^. The tissues were bathed at 37°C, in oxygenated (95% O^2^, 5% CO^2^) Krebs solution (pH 7.4) with 10 mM of glucose and mannitol in the serosal and mucosal compartments, respectively. Tissues were held under voltage-clamp conditions (0 V) and allowed to equilibrate for 15 to 20 min after which basal short-circuit current (Isc, μA/cm^2^) and transepithelial potential were measured in order to calculate basal transepithelial resistance according to Ohm’s law (TER, Ω/cm²). Changes in net electrogenic ion flux were evaluated in the ileum by measuring changes in Isc in response to neuronal depolarization with 10 µM veratridine (Calbiochem, San Diego, CA, USA) or in response to muscarinic receptor stimulation with 100 µM carbachol (Sigma). The difference between basal Isc and peak Isc recorded after veratridine or carbachol addition was measured (ΔIsc, μA/cm^2^). A positive ΔIsc indicated a luminally directed negative net charge transfer (anion secretion). Measurements were conducted and averaged in two adjacent intestinal segments from the same mouse for every gut region (4-5 mice/treatment group).

### DSS Colitis Model

DSS-induced colitis model was performed as previously described^55^, with small modifications. Briefly, F1 mice at 7 weeks of age (7-11/treatment group) were treated for 5 consecutive days with 1.5% DSS (Alfa Aesar, Haverhill, MA, USA) in sterile drinking water. DSS solution was prepared by dissolving the colitogenic agent in distilled water, followed by filter sterilization. DSS consumption was monitored daily by determination of water volume consumed per cage. Health checks and body weight measurements were performed daily to follow disease progression. Feces were obtained for assaying fecal markers of inflammation. Occult blood was determined in Hemoccult Fecal Occult Blood Slide Test System (Beckman Coulter, Brea, CA, USA) and lipocalin-2 (Lcn-2) protein was assayed in Mouse Lipocalin-2/NGAL DuoSet ELISA (R&D Systems, Minneapolis, MN, USA). At the end of DSS treatment, mice were anesthetized with isoflurane and sacrificed by cervical dislocation. Sections of large intestines were obtained for histology and flash-frozen for cytokine measurements in U-Plex T Cell Combo Assay Kit in MESO QuickPlex SQ 120 (Meso Scale Discovery, Rockville, MD, USA). Sections for histology were fixed in Neutral Buffered Formalin 10% (EMD Chemicals, Gibbstown, NJ, USA) and embedded in paraffin. Histological cuts were stained with hematoxylin and eosin for inflammation scoring as described below. Flash frozen tissues were weighed and lysed in 600 μl of lysis buffer, constituted of 150 mM sodium chloride (EMD Chemicals), 20 mM Tris-Hydrochloride pH 7.5 (EMD Chemicals), 1mM EGTA (ethylene glycol-bis(β-aminoethyl ether)-N,N,N′,N′-tetraacetic acid; Sigma), 1% Triton X-100 Surfactant (EMD Chemicals), and 1x cOmplete Protease Inhibitor Cocktail (Roche, Basel, Switzerland). Protein concentration in tissue lysates was determined using the Coomassie (Bradford) Protein Assay Kit (Thermo-Fisher, Waltham, MA, USA).

### Colonic Histologic Scoring

Inflammation was blindly assessed following modified protocol from Chassaing *et al.*^55^ Each section was assigned scores based on the following parameters: (i) inflammatory infiltration (0= none, 1= inflammatory cells above muscularis serosa only, 2= inflammatory cells in submucosa, 3= inflammatory cells in muscularis serosa to muscularis mucosa, 4= extensive inflammation in mucosa and epithelial layer), (ii) crypt damage (0= none, 1= <30%, 2= <60%, 3= only epithelial surface intact, 4= entire crypt and epithelia lost), (iii) goblet cell depletion (0= none, 1= present), and (iv) crypt abscess (0= none, 1= present). Additionally, parameters “i”, “ii” and “iii” were further multiplied by a degree factor of: 1= focal, 2=patchy, or 3=diffuse. Total score ranged from 0-28 per mouse.

## Microbiome Analyses

### DNA Library Preparation

Fecal microbial DNA extracted with DNeasy PowerSoil Pro Kit (Qiagen) was used to amplify the V4 region of the bacterial 16S rRNA gene, and the V3-V4 region of the fungal ITS2 rRNA gene. This generated ready-to-pool dual-indexed amplicon libraries as described previously^56, 57^. 16S amplicon libraries were prepared in house using TopTaq Master Mix kit (Qiagen). Amplicons were cleaned up using QIAquick PCR purification kit (Qiagen), quantified with PicoGreen (Invitrogen, Carlsbad, CA, USA) and diluted to 20 ng/μl for sequencing. ITS2 amplicon libraries were prepared at Microbiome Insights, Vancouver. In house extracted DNA samples were sent to the facility and amplified using Phusion Hot Start II DNA Polymerase (Thermo-Fisher). PCR reactions were cleaned-up and normalized using the high-throughput SequalPrep Normalization Plate Kit (Applied Biosystems) and quantified accurately with the KAPA qPCR Library Quantification kit (Roche). Controls without template DNA, and mock communities with known amounts of selected bacteria and fungi, were included in the PCR and downstream sequencing steps to control for microbial contamination and verify bioinformatics analysis pipeline.

### Sequencing

The pooled and indexed libraries were denatured, diluted, and sequenced in paired-end modus on an Illumina MiSeq (Illumina Inc., San Diego, USA). 16S and ITS rRNA sequencing were performed at Microbiome Insights, UBC. Sequences were checked for quality, trimmed, merged, and checked for chimeras using the DADA2 ^58^ and phyloseq ^59^ packages for R (R Development Core Team; http://www.R-project.org). We built bacterial and fungal community matrices from the resulting unique Amplicon Sequence Variants (ASV).

### Fecal Metabolomics

Metabolites were extracted from gnotobiotic mice fecal pellets by 50% methanol solution. Samples were subjected to untargeted metabolomics analysis using LC-MS. Metabolomics data were acquired by R.A.G. at the Calgary Metabolomics Research Facility, which is supported by the IMC. General metabolomics runs were performed on a Q Exactive HF Hybrid Quadrupole-Orbitrap Mass Spectrometer (Thermo-Fisher) coupled to a Vanquish UHPLC System (Thermo-Fisher). Chromatographic separation was achieved on a Syncronis HILIC UHPLC column (2.1 mm x 100 mm x 1.7 μm, Thermo-Fisher) using a binary solvent system (A+B) at a flow rate of 600 μL/min. Solvent A, 20mM ammonium formate pH 3.0 in mass spectrometry grade H^2^0; Solvent B, mass spectrometry grade acetonitrile (Thermo-Fisher) with 0.1% formic acid (%v/v). A sample injection volume of 2 μL was used. The mass spectrometer was run in negative full scan mode at a resolution of 240 000 scanning from 50-750 m/z.

### Early-Life Immune Assessment

Four-week old F1 mice (5-8/treatment group) were selected for early-life immune assessment. Mice were anaesthetized with isoflurane and blood was obtained by cardiac puncture after confirmed absence of reflex. Following cardiac puncture, mice were sacrificed by cervical dislocation and spleen, jejunum and colon sections were immediately removed for flow cytometry or cytokine measurements as described below. Blood was allowed to clot on ice, spun at 3000 g for 10 min and supernatant (serum) was used to determine immunoglobulin levels. IgE titer was determined in IgE BD OptEIA ELISA (BD Biosciences, San Diego, CA, USA). IgA, IgG1, IgG2a, IgG2b, IgG3 and IgM titers were determined in Mouse Isotyping Panel 1 Assay Kit (Meso Scale Discovery). Jejunum and colon fragments were flash-frozen for cytokine measurements in U-Plex T Cell Combo Assay Kit (Meso Scale Discovery).

### Immunolabeling and Flow Cytometry

Spleens were surgically removed and processed for flow cytometry. Spleens were kept in ice-cold RPMI 1640 + GlutaMax-I medium (Thermo-Fisher) and were macerated in GentleMACS C tubes (Miltenyi Biotec). Cellular macerate was filtered through a 100 μm cell strainer and red blood cells were lysed in 5 ml 1x RBC Lysis Buffer (BioLegend, San Diego, CA, USA). Splenocytes were resuspended in ice-cold cell growth media constituted of RPMI 1640 + GlutaMax-I medium (Thermo-Fisher) supplemented with 10% heat-inactivated Fecal Bovine Serum (Thermo-Fisher), 2000 U Penicillin-Streptomycin (Thermo-Fisher) and 50 μM 2-Mercaptoethanol (Thermo-Fisher). Isolated splenocytes were stained with Fixable Viability Stain (FVS575V; BD Biosciences) and array of intra and extracellular markers for immune cell characterization. Cells were stained for surface markers, fixed, permeabilized with BD Transcription Factor Buffer Set (BD Biosciences) and stained for intracellular markers. The following antibodies were used: PE-Cy7 anti-mouse CD19 (clone 1D3; BD Biosciences), PerCP-Cy5.5 anti-CD11b (M1/70; BD Bioscience), APC-H7 anti-mouse CD8a (53-6.7; BD Bioscience), Alexa Fluor 700 anti-mouse CD3 (17A2; BD Bioscience), V500 anti-mouse CD4 (RM4-5; BD Biosciences), Brilliant Violet 421 anti-mouse CD25 (PC61; BD Biosciences), Brilliant Violet 421 anti-mouse CD11c (HL3; BD Biosciences), PE anti-Gata3 (L50-823; BD Biosciences), PE anti-mouse IL-17A (TC11-18H10; BD Biosciences), PE anti-mouse IL-12/23 (C15.6; BD Biosciences), PE anti-mouse F4/80 (T45-2342; BD Biosciences), Alexa Fluor 488 anti-mouse IL-4 (11B11; BD Biosciences), Alexa Fluor 488 anti-mouse FoxP3 (MF23; BD Biosciences), Alexa Fluor 488 anti-mouse INF-γ (XmG1.2; BD Biosciences), Alexa Fluor 488 anti-mouse I-A/I-E (M5/114.15.2; BD Biosciences), eFluor 660 anti-mouse IL-13 (eBio13A; eBioscience, San Diego, CA), APC anti-mouse IL-10 (JES5-16E3; BD Biosciences), APC anti-mouse IL-6 (MP5-20F3; BD Biosciences), and APC anti-mouse CD103 (M290; BD Biosciences). Samples were run in BD FACSCanto II (BD Biosciences), in the Nicole Perkins Microbial Communities Core Lab, Snyder Institute for Chronic Diseases, University of Calgary.

### OVA-Induced Airway Inflammation Model

Experimental murine OVA model was followed as previously described ^20^ with modifications. F1 mice (5-10/treatment group) were systemically sensitized intraperitoneally (IP) with sterile 200 μg of grade V OVA (Sigma) and 1.3 mg of aluminum hydroxide (Thermo-Fisher) at weeks 7 and 9 post-birth. Following systemic sensitization, airway inflammation was induced by intranasal challenges at week 10 with 50 μg of Low Endo OVA (Whortington, Lakewood, NJ, USA) for three consecutive days, followed by two days of 100 μg of grade V OVA (Sigma) in sterile phosphate-buffered saline (PBS). Intranasal challenges were done in mice under anesthesia with isoflurane. Following challenge scheme, mice were anesthetized with ketamine (200 mg/kg; Vetoquinol, Lavaltrie, QC, Canada) and xylazine (10 mg/kg; Bayer Inc., Mississauga, ON, Canada), and BAL were performed by 3x 1 ml washes with PBS for total and differential cell counts. Total BAL counts were performed in hemocytometer. Differentials (eosinophils, neutrophils, macrophages, and lymphocytes) were performed from 200 cells in hematoxylin and eosin stained CytoSpins (Shandon Cytospin II, Thermo-Shandon, Runcorn, Cheshire, UK) based on standard morphological criteria.

### Data Visualization and Statistical Analysis

We measured gut bacterial alpha-diversity using the Chao1 (richness) and Shannon indices. Non-parametric Mann-Whitney tests were performed to test for significant differences in alpha-diversity between treatments and time points. To account for potential heteroskedasticity in community beta-diversity dispersion and avoid the loss of information through rarefaction^60^, a variance stabilizing transformation was performed prior to any statistical tests^60, 61^. Changes in gut bacterial community structure (beta-diversity) were assessed statistically using Permutational Multivariate Analysis Of Variance (PERMANOVA) ^32^ and visualized using Principal Coordinate Analysis (PCoA) based on Bray-Curtis dissimilarities. To explore further the changes in taxonomical community structure, significant changes in relative abundance of the 9 most dominant bacterial and 8 most dominant fungal ASVs were tested using non-parametric Kruskal-Wallis tests followed by post-hoc Dunn tests with Benjamin-Holmes False Discovery Rate (FDR) correction. Boxplots were made to identify outliers and normality was assessed using the Shapiro-Wilk test. If the data was normally distributed, parametric one-way analysis of variance (ANOVA) with Tukey’s post-hoc tests was used. Correlations between the 30 most abundant 16S and ITS2 ASVs and the detected immunological features in early-life were assessed using bicor method with FDR. We also used SGCCA to understand the correlation structure between different measurements (16S, ITS, immune features) while having the possibility to use dimension reduction especially for the highly correlated metabolomics data. R package mixomics ^62^ was used to perform SGCCA analysis. ITS and 16s data were first changed to ranks ^58^ and all measurements were range normalized. For longitudinal analyses, statistical tests on univariate response variables were performed using a linear mixed-model for repeated measures, followed by an ANOVA with Tukey’s post hoc when appropriate. PCA, ANOVA, T-tests and pathway analysis of fecal metabolites originating from different mice groups was performed by using Metaboanalyst v.4.0^63^. In all tests, significance was set at P<0.05. All data points represent biological replicates. Graphs were made using either R (R Development Core Team) or Prism version 8.1.2 (GraphPad Software, La Jolla, CA, USA). Figures 1, 2, and 6 were created using diagrams from BioRender.com (https://biorender.com/).

## Acknowledgements

The authors would like to acknowledge the following collaborators for their support: Markus Geuking, Simon Hirota and Jens Walter for conceptual feedback and/or help with study design; Dylan Pillay and Alberta Public Laboratories for the fungi isolates; Laurie Wallace, Cristiane Baggio and Kristoff Nieves for technical support; Matheus Heitor and Carolyn Thomson for their help with flow cytometry approaches; Kristy Brown, Ian Lewis and Ryan Groves for metabolomics support. This work was supported by funds from the Cumming School of Medicine, The Alberta Children Hospital Research Institute, the Snyder Institute of Chronic Diseases, Canadian Institutes for Health Research, Sick Kids Foundation, and the Canadian Lung Association. K.A.S. holds the Crohn’s and Colitis Canada Chair in IBD Research. E.V.T.B. is funded by the Eyes High Doctoral Recruitment Scholarship. N.G.J. is funded by the Parker B Francis Fellowship. J.-B.C. is funded by the Human Frontier Science Program. F.A.V. is funded by the National Council for Scientific and Technological Development (CNPq/Brazil). The International Microbiome Centre (IMC) is supported by the Cumming School of Medicine, University of Calgary, Western Economic Diversification (WED) and Alberta Economic Development and Trade (AEDT), Canada.

## Author contributions

E.V.T.B., V.K.P., M.W.G., I.L.L., K.M., W.K.M., K.A.S., M.-C.A. contributed to study design; E.V.T.B., V.K.P., M.W.G., I.L.L., M.-C.A. carried out the mouse experiments; E.V.T.B., V.K.P., M.W.G., I.L.L., N.G.J., J.-B.C., F.A.V., C.M.K., J.S., M.-C.A. prepared all the samples; E.V.T.B., M.W.G., M.-C.A. optimized the qPCR strategy; M.W.G., M.-C.A. performed the qPCR analysis; J.-B.C., F.A.V., C.M.K. performed and analyzed gut physiology experiments; E.V.T.B., M.W.G., M.-C.A. analyzed DSS-colitis data; E.V.T.B., I.L.L., J.S., M.-C.A. prepared libraries and optimized sequencing strategy; E.V.T.B., I.L.L., M.-C.A. analyzed the 16S and ITS data; V.K.P., M.-C.A. analyzed the metabolomics data; E.V.T.B., M.-C.A. analyzed the early-life immunity data; E.V.T.B., M.W.G., N.G.J., R.J.A.W., M.M.K. analyzed airway inflammation data; E.V.T.B., V.K.P., M.W.G., I.L.L., M.-C.A. curated all the metadata; H.R. performed SGCCA for multi-omics correlation; E.V.T.B., M.-C.A. wrote the manuscript; E.V.T.B., M.W.G., V.K.P., N.G.J., K.D.M., C.M.K, F.A.V., J.-B. C., K.A.S., M.-C.A. edited the manuscript. All authors contributed extensively to the work presented in this paper

## Data and materials availability

16S and ITS sequence reads were deposited to European Nucleotide Archive (ENA, https://www.ebi.ac.uk/ena).

## Supplementary Materials

Supplementary Fig. 1: Fungi colonize mouse intestinal tract less efficiently than bacteria.

Supplementary Fig. 2: Microbial colonization does not impact intestinal transit and colonic functionality.

Supplementary Fig. 3: DSS-induced colitis in gnotobiotic mice.

Supplementary Fig. 4: Fungal colonization and antimicrobial treatments impact microbial diversity and community composition.

Supplementary Fig. 5: Metabolic profiles of small intestine content is not different among gnotobiotic groups.

Supplementary Fig. 6 Flow cytometry gating strategy.

Supplementary Fig. 7: Additional differences reported in systemic immunity.

Supplementary Fig. 8: Fungal colonization impacts intestinal immunity.

Supplementary Fig. 9: Network plot for microbial ASVs and immune features.

Supplementary Table 1: Strains and primers used in this study.

Supplementary Table 2: Permutational multivariate analysis of mice gut bacterial (16S) taxonomic community structure.

Supplementary Table 3: Permutational multivariate analysis of mice gut fungal (ITS2) taxonomic community structure.

Supplementary Table 4: Metabolites with statistically significant differences detected among gnotobiotic groups.

Supplementary Table 5: Metabolites with statistically significant differences detected between Y and GF gnotobiotic groups.

