## Supplementary material for "Contribution of fungal microbiome to intestinal physiology, early-life immune development and mucosal inflammation in mice"

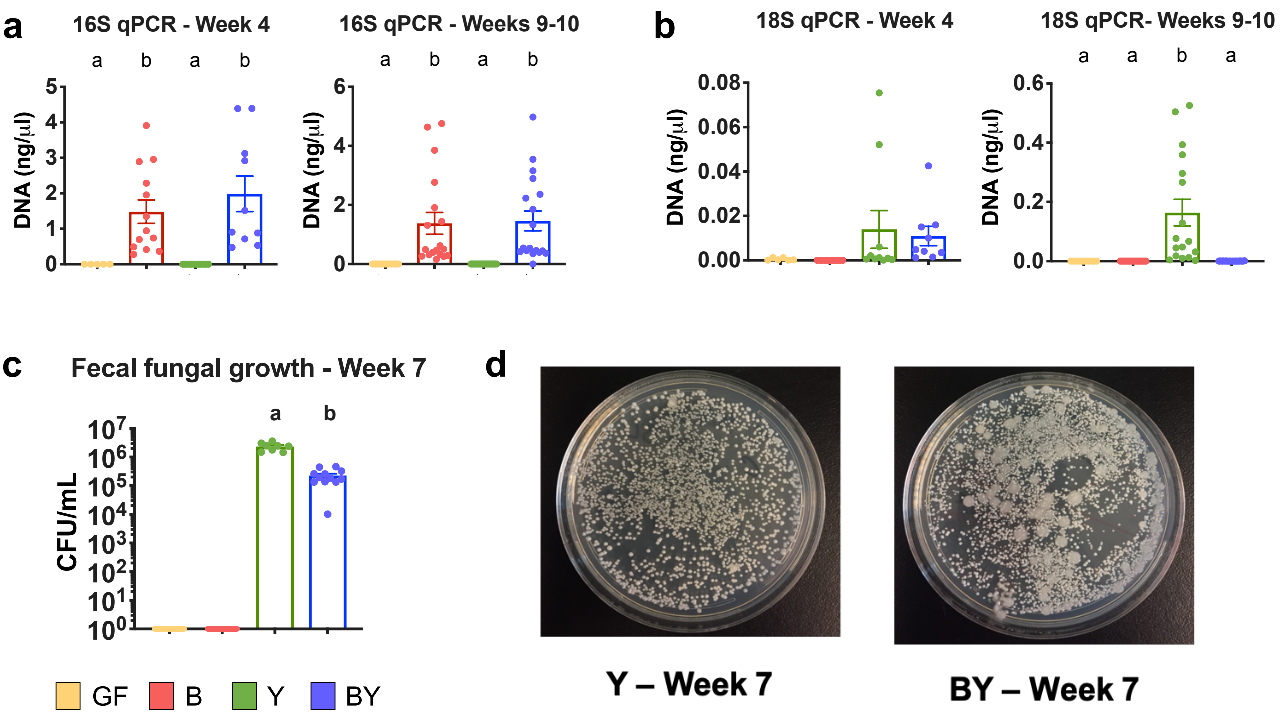


**Supplementary Fig. 1: Fungi colonize mouse intestinal tract less efficiently than bacteria.** qPCR quantification (standard curve method) of **(a)** bacterial and **(b)** fungal DNA in fecal samples by amplification of the 16S and 18S rRNA genes, respectively. **(c)** Fecal fungal colony counts in YM agar media supplemented with antibiotics (gentamycin + chloramphenicol). **(d)** Fecal fungal growth in selective medium from representative Y and BY mice. (a-c) Color denotes colonization treatment (GF = yellow, B = red, Y = green, BY = royal blue); (a-b) Data combined from two different experiments, N=5-19; (c) N=6-12; different letters above bars indicate statistically significant differences defined by ANOVA and Tukey posthoc tests; P<0.05.

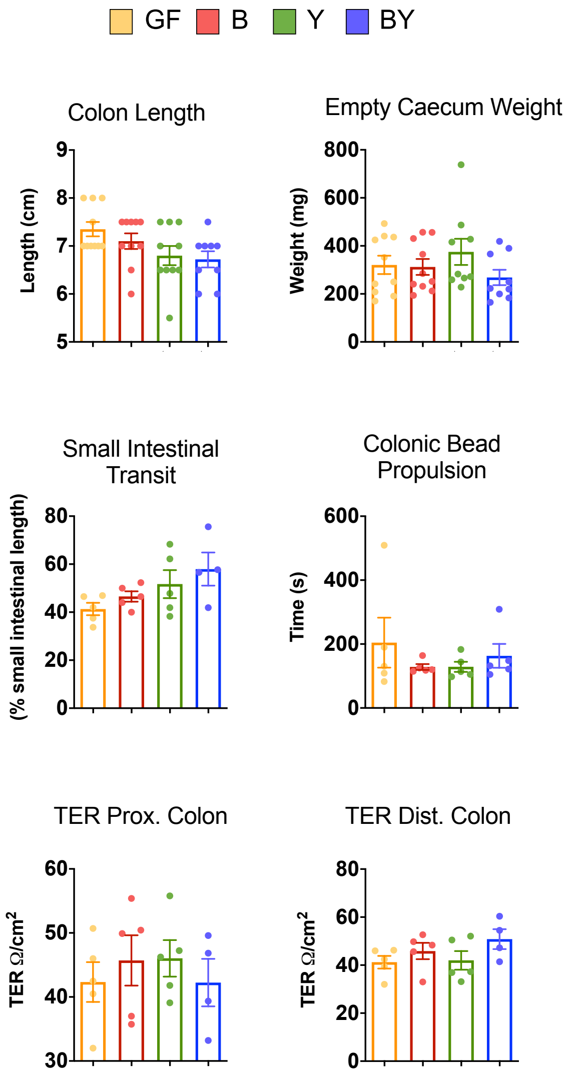


**Supplementary Fig. 2: Microbial colonization does not impact intestinal transit and colonic functionality. (a)** Large intestine length and empty caecum weight of gnotobiotic dams. **(b)** Small and large intestine transit determined by movement of non-absorbable dye and colonic bead propulsion, respectively. **(c)** Proximal and distal colon paracellular permeability measured by electrical Transepithelial Resistance (TER). (a-c) Color denotes colonization treatment (GF = yellow, B = red, Y = green, BY = royal blue); (a) N=8-10; (b-c) N=4-5; no statistically significant differences between groups (ANOVA and Tukey posthoc tests).

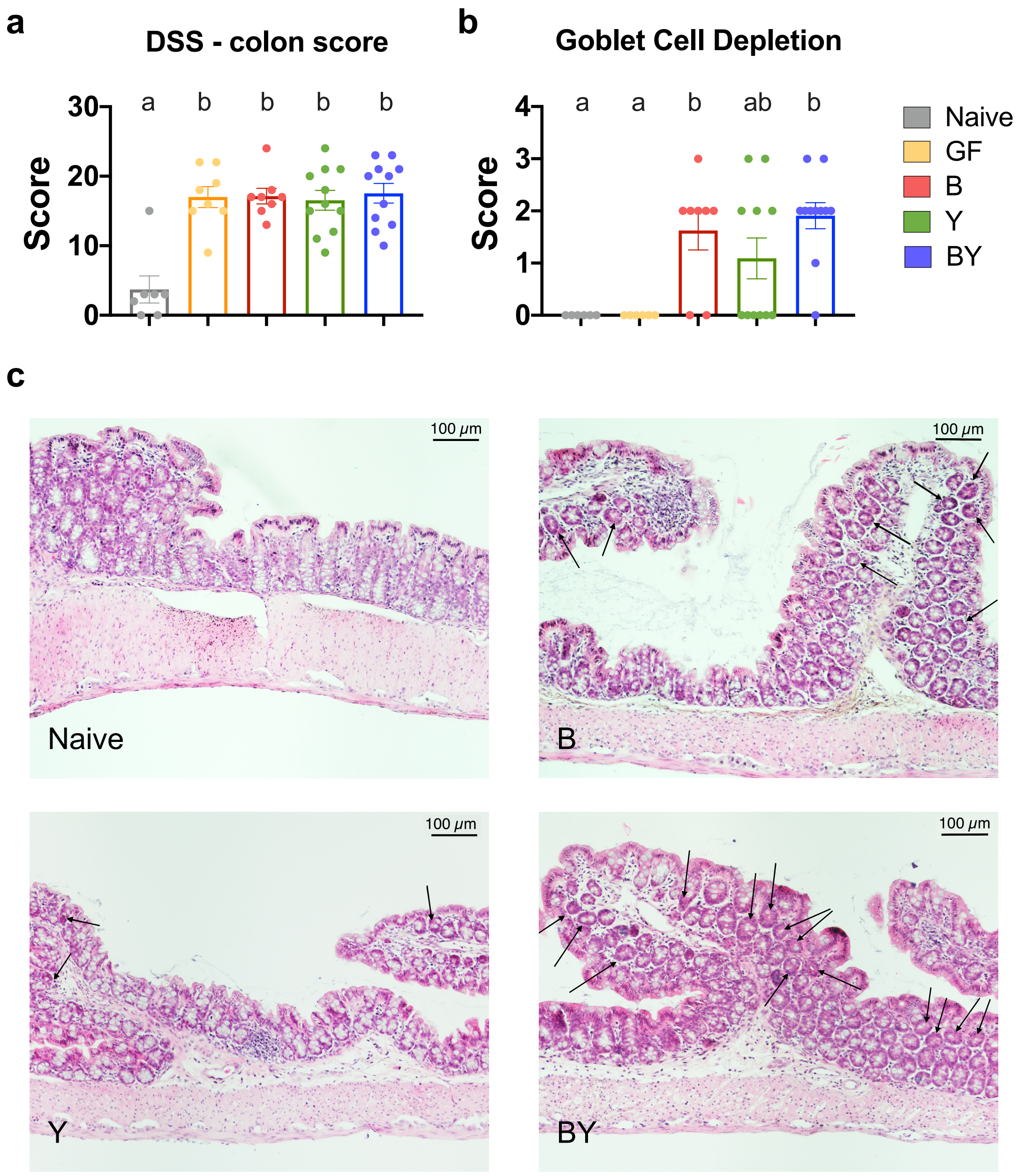


**Supplementary Fig. 3: DSS-induced colitis in gnotobiotic mice. (a)** Total histopathological inflammation score for DSS-challenged colon. Inflammation score was assessed by sum of pathological measurements, including total inflammation, crypt damage, crypt abscess and goblet cell depletion. **(b)** Results for goblet cell depletion. Presence (1) or absence (0) of goblet cell depletion was multiplied by degree score focal (1), patchy (2) or diffuse (3). (a-b) Color denotes colonization and DSS treatment (Naive = gray, GF = yellow, B = red, Y = green, BY = royal blue); N=7-11; different letters above bars indicate statistically significant differences defined by ANOVA and Tukey posthoc tests; $P<0.05$. **(c)** Representative hematoxylin and eosin (H&E)-stained sections for gnotobiotic groups. Arrows indicate regions of goblet cell depletion. Scale bar, 100 μm.

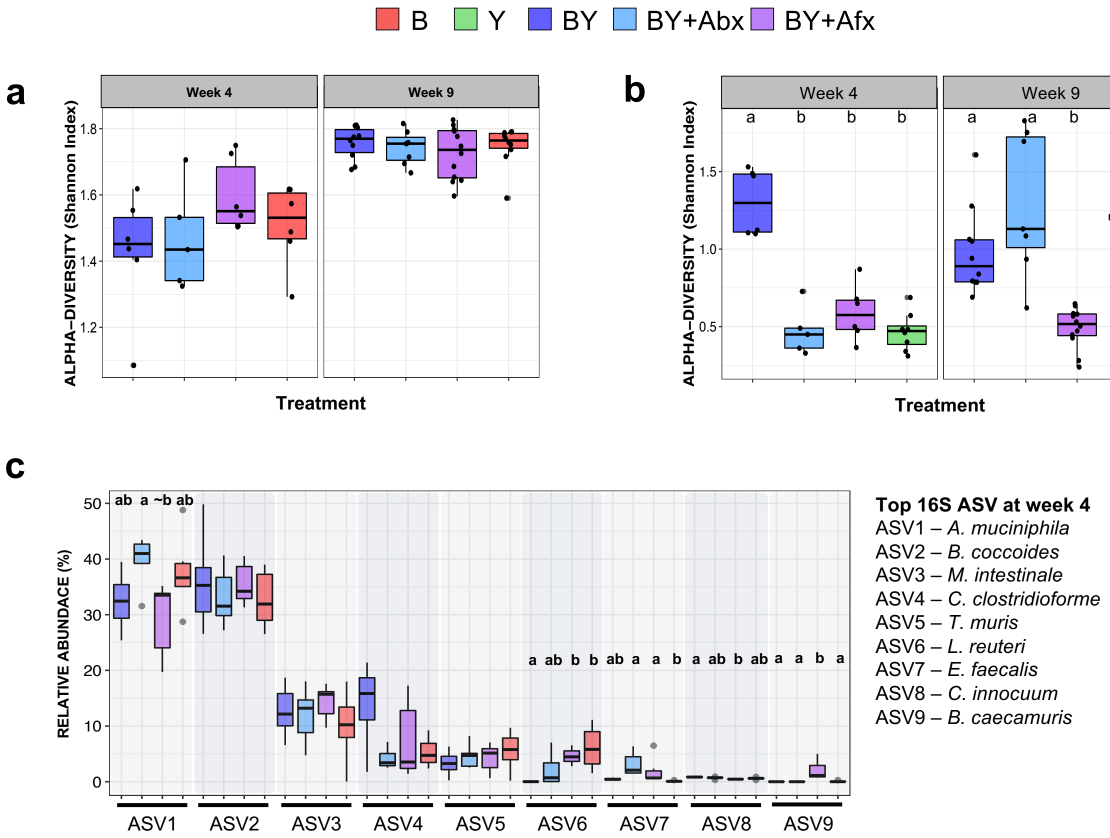


**Supplementary Fig. 4: Fungal colonization and antimicrobial treatments impact microbial diversity and community composition.** Ecological community analyses of 16S and ITS2 rRNA sequences. Plots of **(a)**16S and **(b)** ITS2 alpha-diversity (Shannon Index). **(c)** Relative abundances of the 9 most dominant bacterial ASVs at 4 weeks. (a-c) Color denotes colonization treatment (B = red, Y = green, BY = royal blue, BY+Abx = cyan blue, BY+Afx = purple); $N_{week4}$=5-8; $N_{week9}$=7-12; different letters above bars indicate statistically significant differences defined by Kruskal-Wallis with post-hoc Dunn tests and FDR corrected; P<0.05.

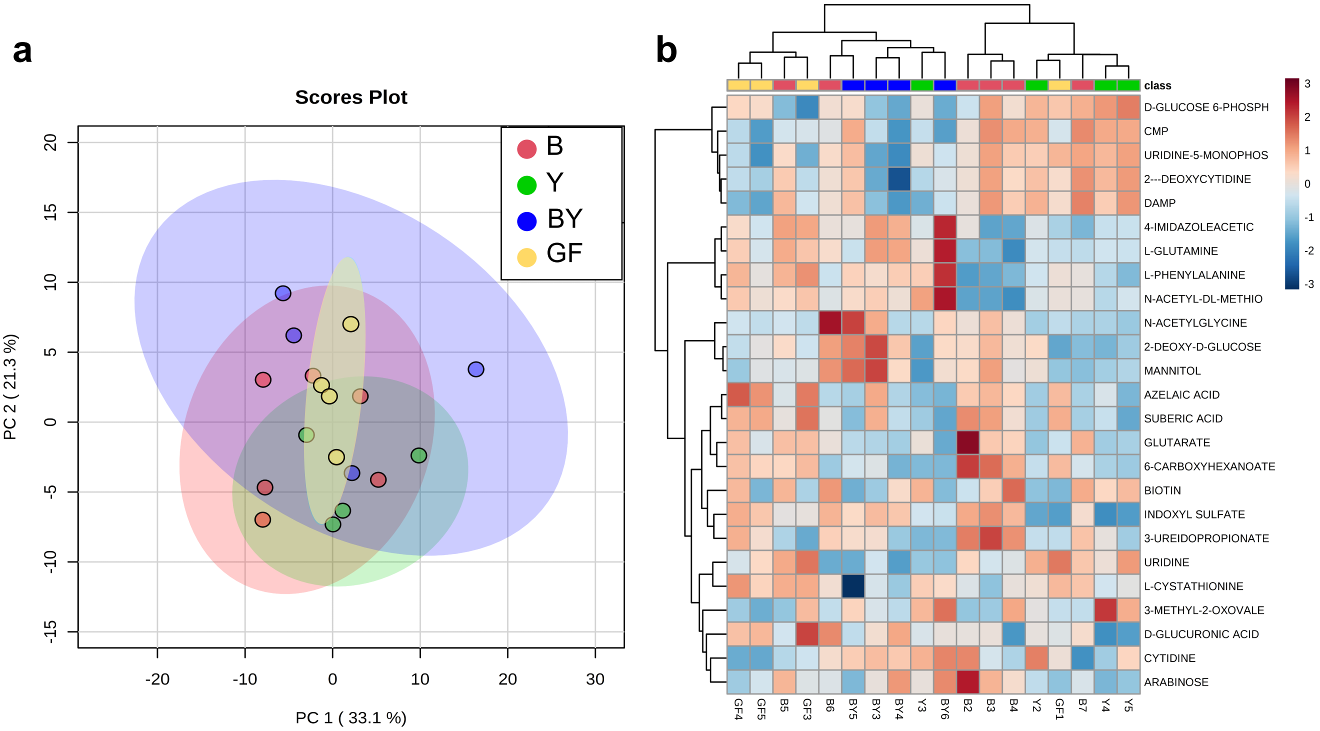


**Supplementary Fig. 5: Metabolic profiles of small intestine content is not different among gnotobiotic groups. (a)** Principal component analysis score plot of 124 metabolites detected in small bowel fecal content of gnotobiotic mice groups at 4 weeks of age. **(b)** Heat map of expression of top 25 metabolites detected in small bowel contents at 4 weeks. (a-b) Color denotes colonization treatment (GF = yellow, B = red, Y = green, BY = royal blue); N=4-6.

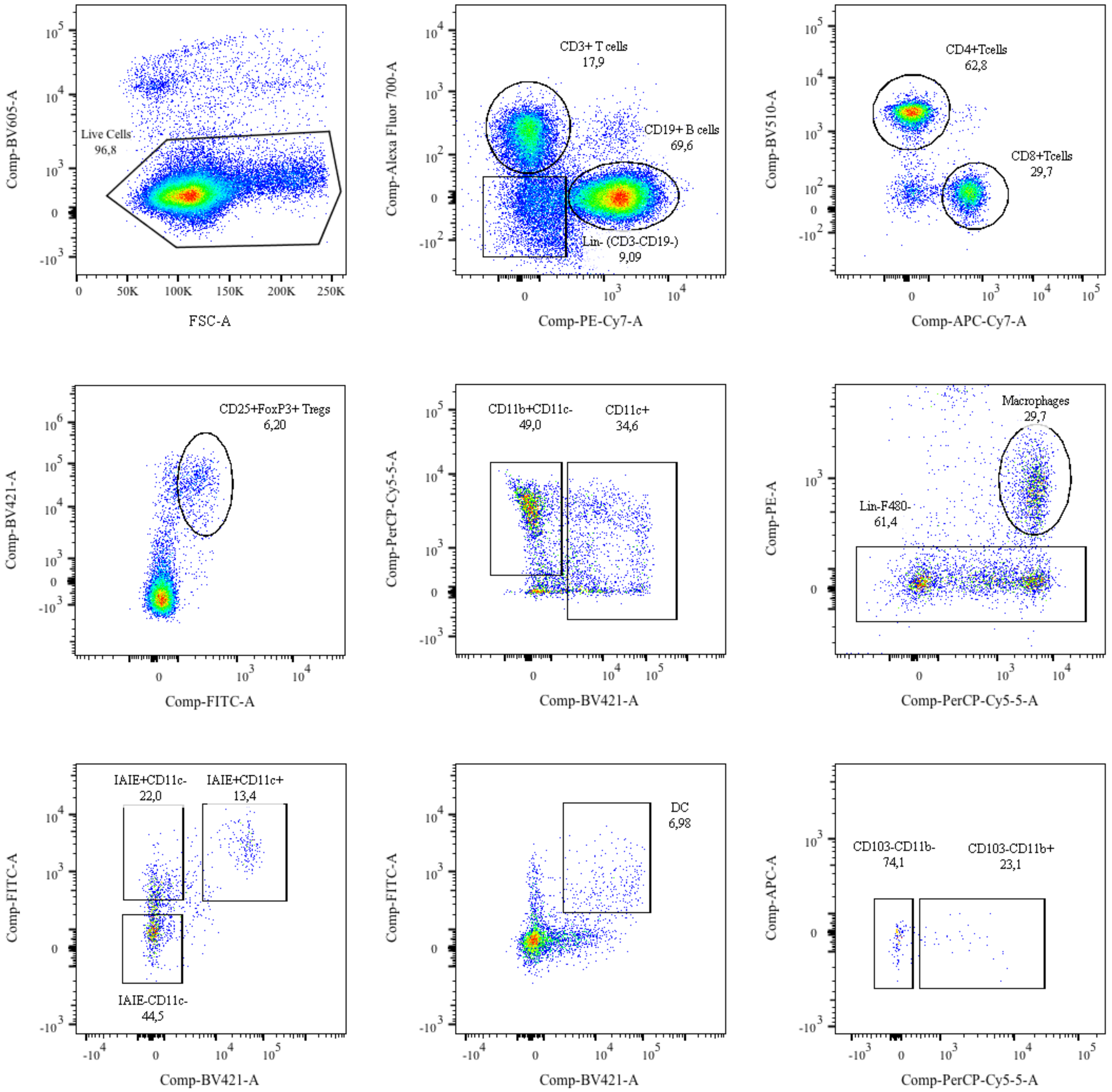


**Supplementary Fig. 6: Flow cytometry gating strategy.** Spleen immune cells were stained with intra- and extracellular marker-specific antibodies and sorted by flow cytometry (see methods). Analysis performed in FlowJo$^{TM}$ version 10.5.3.

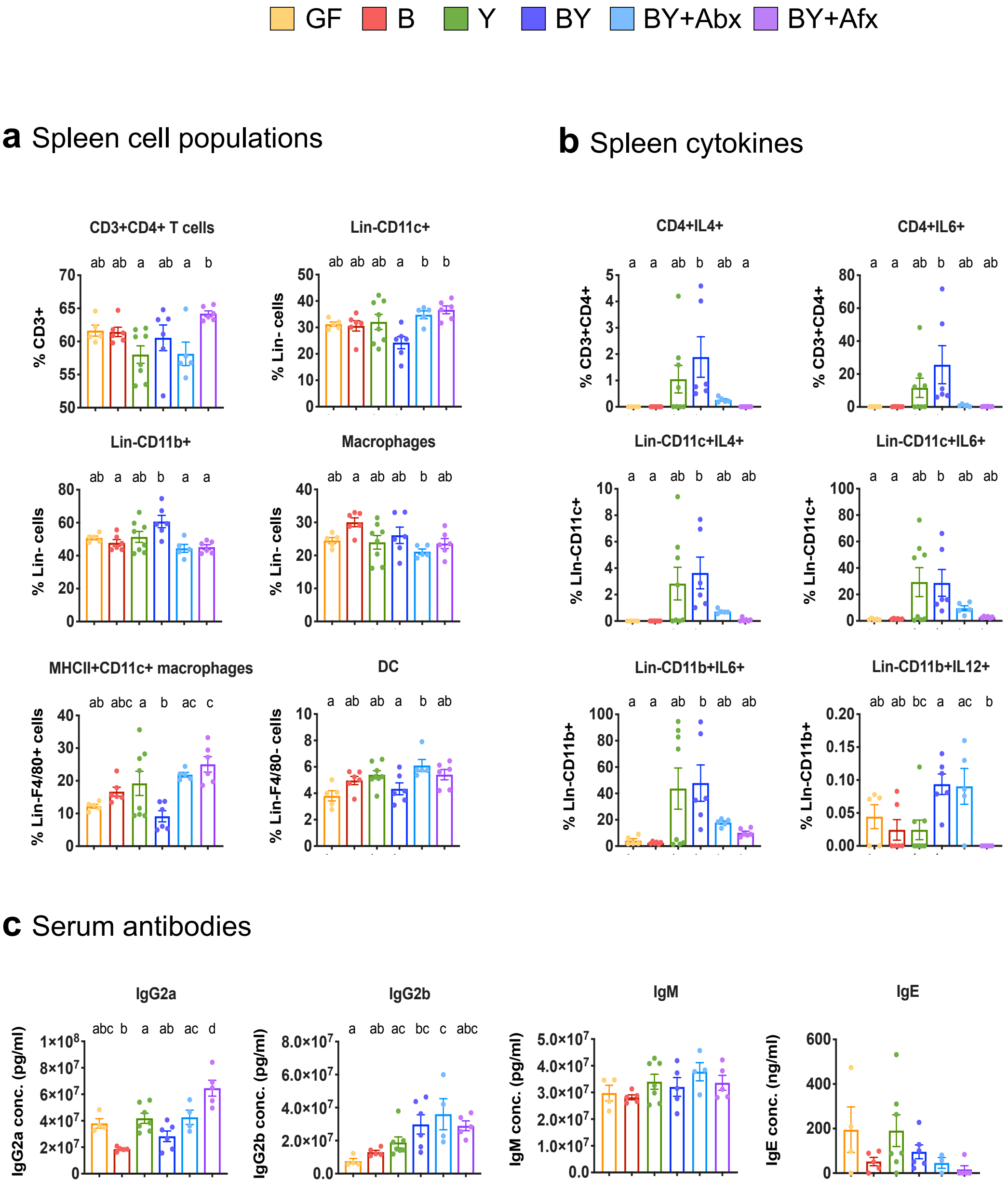


**Supplementary Fig. 7: Additional differences reported in systemic immunity.** Percentage of **(a)** spleen cell populations and **(b)** cytokine-producing splenocytes from 4-week-old gnotobiotic mice. **(c)** Serum antibody concentrations detected by electrochemiluminescence (MSD – IgG2a, IgG2b and IgM) or ELISA (IgE). (a-c) Color denotes colonization treatment (GF = yellow, B = red, Y = green, BY = royal blue, BY+Abx = cyan blue, BY+Afx = purple); N= 5-8; different letters above bars indicate statistically significant differences defined by ANOVA and Tukey posthoc tests; $P<0.05$.

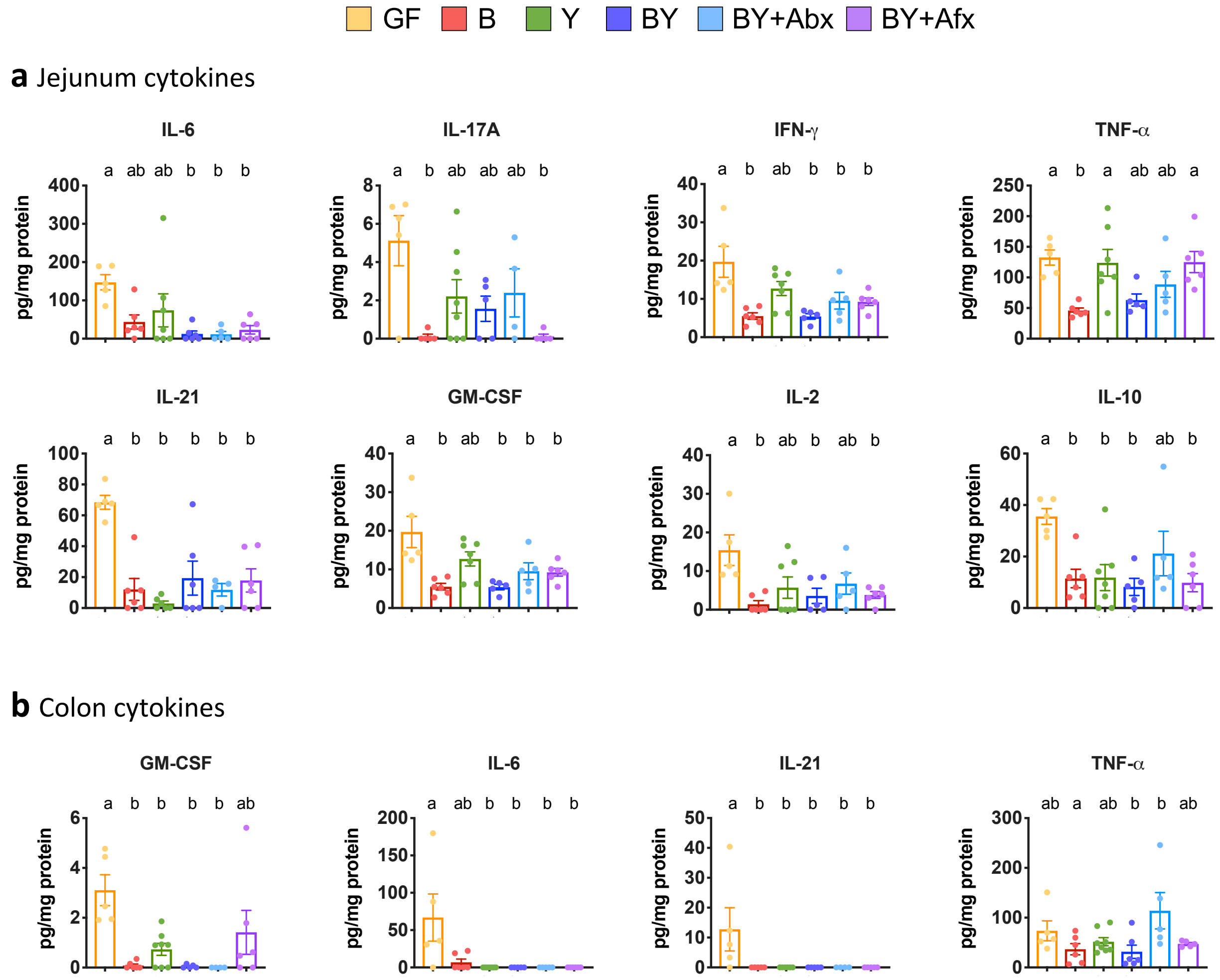


**Supplementary Fig. 8: Fungal colonization impacts intestinal immunity.** Cytokine concentration in **(a)** jejunum and **(b)** colon lysates detected by electrochemiluminescence (MSD). (a-b) Color denotes colonization treatment (GF = yellow, B = red, Y = green, BY = royal blue, BY+Abx = cyan blue, BY+Afx = purple); N= 5-8; different letters above bars indicate statistically significant differences defined by ANOVA and Tukey posthoc tests; $P<0.05$.

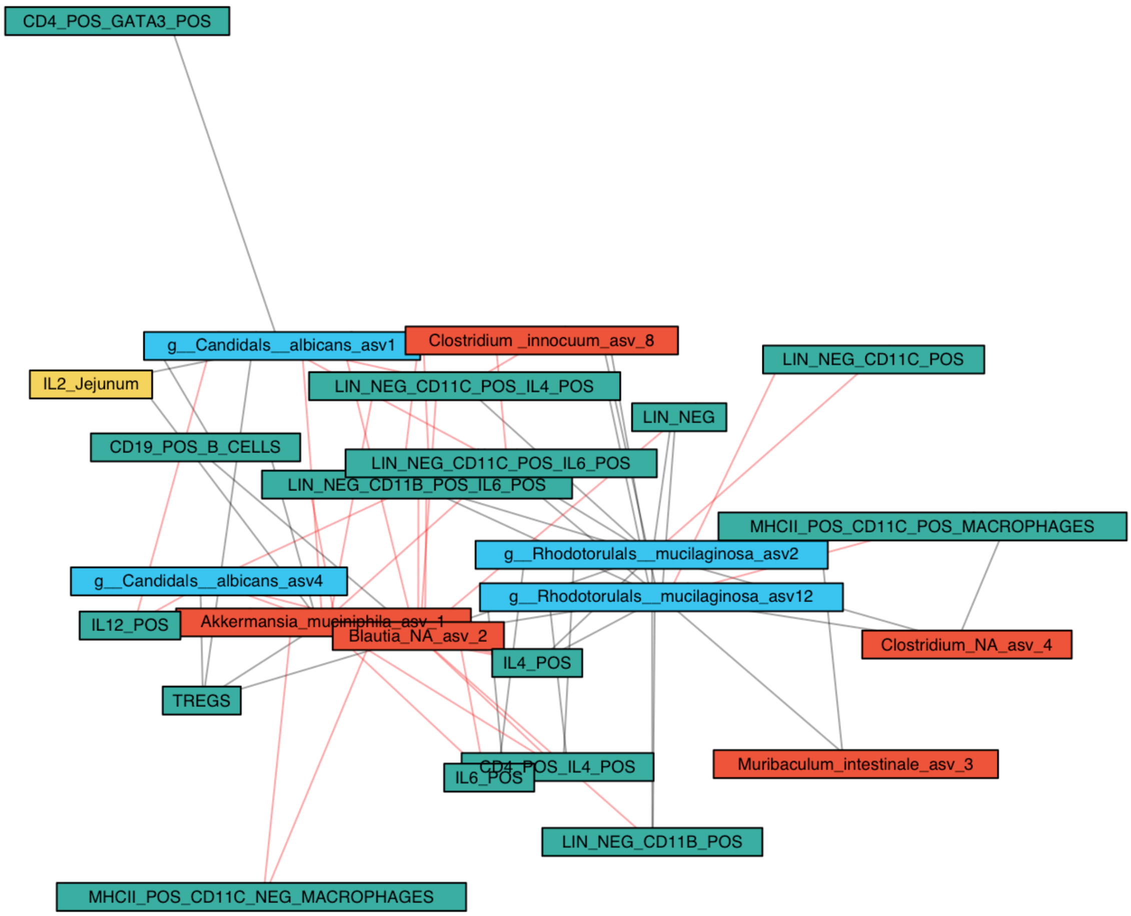


**Supplementary Fig. 9: Network plot for microbial ASVs and immune features.** Relevance network plots for bacteria (Persian red) and fungi (blue) ASVs with the systemic (green) and intestinal (yellow) immune features. Relevant correlations are denoted with a line (red=positive, black=negative) and defined by pair-wise similarity matrix for SGCCA.

**Supplementary Table 1: Strains and primers used in this study.**

| sDMDMm2 Consortia | Source | References |
|---|---|---|
| '*Acutalibacter muris*' – KB18 | Dr. K. D. McCoy | 1 |
| *Akkermansia muciniphila* – YL44 | Dr. K. D. McCoy | 1 |
| '*Bacteroides caecimuris*' – I48 | Dr. K. D. McCoy | 1 |
| *Blautia coccoides* – YL58 | Dr. K. D. McCoy | 1 |
| *Clostridium clostridioforme* – YL32 | Dr. K. D. McCoy | 1 |
| *C. innocuum* – I46 | Dr. K. D. McCoy | 1 |
| *Enterococcus faecalis* – KB1 | Dr. K. D. McCoy | 1 |
| *Flavonifractor plautii* – YL31 | Dr. K. D. McCoy | 1 |
| *Lactobacillus reuteri* – I49 | Dr. K. D. McCoy | 1 |
| '*Turicimonas muris*' – YL45 | Dr. K. D. McCoy | 1 |
| *Muribaculum intestinale* – YL27 | Dr. K. D. McCoy | 1 |
| *Bifidobacterium longum* subsp. *animalis* – YL2 | Dr. K. D. McCoy | 1 |
| **Fungal Strains** | | |
| *Candida albicans* | Dr. D. R. Pillai/APL | |
| *C. glabrata* | Dr. D. R. Pillai/APL | |
| *C. krusei (Issatchenkia orientalis)* | Dr. D. R. Pillai/APL | |
| *C. parapsilosis* | Dr. D. R. Pillai/APL | |
| *Pichia kudriavzevii* | DSMZ | |
| *Rhodotorula mucilagin*osa | DSMZ | |
| **Universal 16S primers** | | |
| U16Sfr: 5'-TCC TAC GGG AGG CAG CAG T-3' | | 2 |
| U16Srv: 5'- GGA CTA CCA GGG TAT CTA ATC CTG TT-3' | | 2 |
| **Fungal 18S primers** | | |
| FR1: 5′-AIC CAT TCA ATC GGT AIT-3′ | | 3 |
| FF390: 5′-CGA TAA CGA ACG AGA CCT-3′ | | 3 |

**Abbreviations:**
APL = Alberta Public Laboratories; DSMZ = Leibniz Institute DSMZ-German Collection of Microorganisms and Cell Cultures GmbH.

**Supplementary Table 2**: **Permutational multivariate analysis of mice gut bacterial (16S) taxonomic community structure.** Permutational multivariate analysis of variance stabilizing transformed community matrix testing the influence of cage ID, treatment type and collection week.

| **Whole dataset** | | **16S Permutation** | | |
|---|---|---|---|---|
| **Variable** | DF | F.Model | $R^2$ (%) | P-Value* |
| **Treatment** | 3 | 3.2603 | 14.43 | <0.001 |
| **Collection week** | 1 | 12.085 | 16.76 | <0.001 |
| **Week 4** | | | | |
| **Variable** | DF | F.Model | $R^2$ (%) | P-Value* |
| **Treatment** | 3 | 7.1551 | 51.72 | <0.001 |
| **Cage ID** | 1 | 2.0340 | 4.90 | 0.097 |
| **Week 9** | | | | |
| **Variable** | DF | F.Model | $R^2$ (%) | P-Value* |
| **Treatment** | 3 | 2.5788 | 16.37 | 0.002 |
| **Cage ID** | 7 | 1.6452 | 24.37 | 0.006 |

**Supplementary Table 3: Permutational multivariate analysis of mice gut fungal (ITS2) taxonomic community structure.** Permutational multivariate analysis of variance stabilizing transformed community matrix testing the influence of cage ID, treatment type and collection week.

| **Whole dataset** | | **ITS2 Permutation** | | |
|---|---|---|---|---|
| **Variable** | DF | F.Model | $R^2$ (%) | P-Value* |
| **Treatment** | 3 | 14.308 | 39.87 | <0.001 |
| **Collection week** | 1 | 4.721 | 4.38 | 0.002 |
| **Week 4** | | | | |
| **Variable** | DF | F.Model | $R^2$ (%) | P-Value* |
| **Treatment** | 3 | 13.175 | 63.98 | <0.001 |
| **Cage ID** | 2 | 1.624 | 5.25 | 0.093 |
| **Week 9** | | | | |
| **Variable** | DF | F.Model | $R^2$ (%) | P-Value* |
| **Treatment** | 3 | 19.7925 | 54.45 | <0.001 |
| **Cage ID** | 7 | 2.9524 | 18.95 | <0.001 |

* P-values of statistical significance defined by PERMANOVA on Bray-Curtis distances.

**Supplementary Table 4: Metabolites with statistically significant differences detected among gnotobiotic groups.** List of 99 metabolites with statistically significant differences in relative abundance among gnotobiotic groups. Metabolites in fecal samples were extracted in 50% methanol solution and quantified by LC-MS in a Q Exactive HF Hybrid Quadrupole-Orbitrap Mass Spectrometer coupled to a Vanquish UHPLC System (see methods). Statistical significant differences determined by ANOVA with Fisher's post-hoc + FDR; P<0.05.

| Compound | F-Value | P-Value | -Log10(P) | FDR |
|---|---|---|---|---|
| SHIKIMATE | 712.72 | 3.25E-30 | 29.4890 | 4.38E-28 |
| L-HISTIDINE | 127.89 | 2.73E-19 | 18.5640 | 1.79E-17 |
| L-ASPARAGINE | 124.56 | 3.97E-19 | 18.4010 | 1.79E-17 |
| SUCROSE | 111.99 | 1.80E-18 | 17.7450 | 6.07E-17 |
| 5-HYDROXY-L-TRYPTOPHAN | 97.022 | 1.36E-17 | 16.8660 | 3.68E-16 |
| 3-SULFINO-L-ALANINE | 94.462 | 1.98E-17 | 16.7030 | 4.46E-16 |
| 6-HYDROXYNICOTINATE | 90.622 | 3.54E-17 | 16.4500 | 6.84E-16 |
| THYMIDINE | 87.529 | 5.76E-17 | 16.2400 | 9.71E-16 |
| N-formyl-L-methionine | 86.743 | 6.53E-17 | 16.1850 | 9.79E-16 |
| NICOTINATE | 84.825 | 8.91E-17 | 16.0500 | 1.20E-15 |
| GUANOSINE | 80.509 | 1.84E-16 | 15.7350 | 2.26E-15 |
| N-ACETYL-L-PHENYLALANINE | 73.039 | 7.06E-16 | 15.1510 | 7.95E-15 |
| N-ACETYL-L-ALANINE | 71.962 | 8.66E-16 | 15.0620 | 8.46E-15 |
| L- Carnosine | 71.895 | 8.78E-16 | 15.0570 | 8.46E-15 |
| STACHYOSE | 62.985 | 5.35E-15 | 14.2720 | 4.70E-14 |
| 5-OXO-D-PROLINE | 62.797 | 5.57E-15 | 14.2540 | 4.70E-14 |
| DEOXYCYTIDINE | 62.437 | 6.02E-15 | 14.2200 | 4.78E-14 |
| 3-METHYL-2-OXOVALERIC ACID | 58.466 | 1.46E-14 | 13.8350 | 1.10E-13 |
| D-GLUCOSAMINE 6-PHOSPHATE | 54.791 | 3.50E-14 | 13.4560 | 2.49E-13 |
| URATE | 51.999 | 7.03E-14 | 13.1530 | 4.55E-13 |
| 4-METHYL-2-OXO-PENTANOIC ACID | 51.969 | 7.08E-14 | 13.1500 | 4.55E-13 |
| D--RAFFINOSE | 51.632 | 7.73E-14 | 13.1120 | 4.74E-13 |
| L-ASPARTATE | 50.066 | 1.16E-13 | 12.9350 | 6.82E-13 |
| ADENINE | 48.540 | 1.75E-13 | 12.7570 | 9.85E-13 |
| L-GLUTAMINE | 41.542 | 1.34E-12 | 11.8730 | 7.23E-12 |
| MANNITOL | 39.503 | 2.56E-12 | 11.5920 | 1.33E-11 |
| L-SERINE | 36.829 | 6.27E-12 | 11.2030 | 3.13E-11 |
| L-LYSINE | 36.149 | 7.94E-12 | 11.1000 | 3.83E-11 |
| 4-HYDROXYBENZOATE | 35.757 | 9.11E-12 | 11.0400 | 4.19E-11 |
| N-ACETYL-DL-GLUTAMIC ACID | 35.698 | 9.30E-12 | 11.0310 | 4.19E-11 |
| THYMINE | 34.790 | 1.29E-11 | 10.8900 | 5.61E-11 |
| N-ALPHA-ACETYL-L-ASPARAGINE | 32.512 | 3.01E-11 | 10.5220 | 1.27E-10 |
| ITACONATE | 31.191 | 5.03E-11 | 10.2990 | 2.06E-10 |
| L-ORNITHINE | 28.390 | 1.59E-10 | 9.79810 | 6.32E-10 |
| L-ARGININE | 28.185 | 1.74E-10 | 9.75990 | 6.71E-10 |
| XANTHOSINE | 28.062 | 1.83E-10 | 9.73690 | 6.87E-10 |
| TAURINE | 27.225 | 2.64E-10 | 9.57810 | 9.64E-10 |
| 4-HYDROXY-2-QUINOLINECARBOXYLIC ACID | 25.413 | 6.02E-10 | 9.22040 | 2.14E-09 |
| SUCCINATE | 24.363 | 9.91E-10 | 9.00380 | 3.43E-09 |
| L-ARABITOL | 22.336 | 2.73E-09 | 8.56440 | 9.20E-09 |
| 5---DEOXYADENOSINE | 21.599 | 4.01E-09 | 8.39710 | 1.32E-08 |
| N-acetyl-glutamine | 19.308 | 1.42E-08 | 7.84780 | 4.56E-08 |
| L-CYSTEINE | 18.512 | 2.26E-08 | 7.64560 | 7.10E-08 |
| CYTIDINE | 17.608 | 3.91E-08 | 7.40840 | 1.20E-07 |
| URACIL | 16.126 | 1.00E-07 | 6.99990 | 3.00E-07 |
| BIOTIN | 15.928 | 1.14E-07 | 6.94320 | 3.34E-07 |
| URIDINE | 15.145 | 1.93E-07 | 6.71460 | 5.54E-07 |
| DEOXYURIDINE | 14.147 | 3.88E-07 | 6.41140 | 1.09E-06 |
| INOSINE | 14.122 | 3.95E-07 | 6.40370 | 1.09E-06 |
| NALPHA-ACETYL-L-LYSINE | 13.628 | 5.65E-07 | 6.2482 | 1.52E-06 |
| 2---DEOXYADENOSINE | 12.995 | 9.05E-07 | 6.0436 | 2.39E-06 |
| N-ACETYL-L-LEUCINE | 12.766 | 1.08E-06 | 5.9676 | 2.77E-06 |
| S-CARBOXYMETHYL-L-CYSTEINE | 12.755 | 1.09E-06 | 5.9642 | 2.77E-06 |
| XANTHINE | 11.142 | 3.94E-06 | 5.4050 | 9.84E-06 |
| GLUCOSAMINATE | 10.722 | 5.61E-06 | 5.2514 | 1.38E-05 |
| L-CYSTATHIONINE | 10.501 | 6.77E-06 | 5.1693 | 1.63E-05 |
| N-ACETYL-D-TRYPTOPHAN | 9.8940 | 1.15E-05 | 4.9381 | 2.73E-05 |
| L-THREONINE | 9.4238 | 1.76E-05 | 4.7534 | 4.11E-05 |
| N-acetyl-L-threonine | 8.8196 | 3.10E-05 | 4.5085 | 7.07E-05 |
| N-METHYL-D-ASPARTIC ACID | 8.8062 | 3.14E-05 | 4.5029 | 7.07E-05 |
| PTERIN | 7.7336 | 9.02E-05 | 4.0449 | 0.00019956 |
| D-GLUCURONIC ACID | 7.5490 | 0.00010890 | 3.9630 | 0.00023713 |
| SN-GLYCEROL 3-PHOSPHATE | 7.1176 | 0.00017080 | 3.7675 | 0.00036600 |
| FUMARATE | 7.0042 | 0.00019265 | 3.7152 | 0.00040638 |
| CREATINE | 6.9724 | 0.00019930 | 3.7005 | 0.00041392 |
| 4-IMIDAZOLEACETIC ACID | 6.9282 | 0.00020895 | 3.6800 | 0.00042739 |
| L-ALANINE | 6.8943 | 0.00021668 | 3.6642 | 0.00043659 |
| D-GLUCOSE 6-PHOSPHATE | 6.8636 | 0.00022397 | 3.6498 | 0.00044464 |
| N-ACETYL-DL-SERINE | 6.7970 | 0.00024062 | 3.6187 | 0.00047077 |
| DOCOSAHEXAENOIC ACID | 6.5938 | 0.00030014 | 3.5227 | 0.00057292 |
| GUANINE | 6.5903 | 0.00030131 | 3.5210 | 0.00057292 |
| DL-5-HYDROXYLYSINE | 6.5541 | 0.00031351 | 3.5038 | 0.00058782 |
| XANTHURENIC ACID | 6.4717 | 0.00034331 | 3.4643 | 0.00063488 |
| L-TYROSINE | 6.2399 | 0.00044433 | 3.3523 | 0.00080095 |
| MALEIC ACID | 6.2386 | 0.00044497 | 3.3517 | 0.00080095 |
| 3-METHYLADENINE | 5.9952 | 0.00058598 | 3.2321 | 0.00104090 |
| OPHTHALMIC ACID | 5.8025 | 0.00073105 | 3.1361 | 0.00128170 |
| 3-UREIDOPROPIONATE | 5.6646 | 0.00085799 | 3.0665 | 0.00148480 |
| THYMIDINE 5---MONOPHOSPHATE | 5.6538 | 0.00086889 | 3.0610 | 0.00148480 |
| L-CYSTEIC ACID | 5.5149 | 0.0010226 | 2.9903 | 0.00170690 |
| L-LEUCINE | 5.5137 | 0.0010241 | 2.9896 | 0.00170690 |
| SUBERIC ACID | 5.4937 | 0.0010485 | 2.9794 | 0.00172620 |
| L-GLUTAMIC ACID | 5.4328 | 0.0011268 | 2.9481 | 0.00183280 |
| 2-METHYLMALEATE | 4.9243 | 0.0020812 | 2.6817 | 0.00334470 |
| L-PHENYLALANINE | 4.7888 | 0.0024600 | 2.6091 | 0.00390710 |
| HYPOXANTHINE | 4.7512 | 0.0025775 | 2.5888 | 0.00402760 |
| DEHYDROASCORBATE | 4.7456 | 0.0025956 | 2.5858 | 0.00402760 |
| D-PANTOTHENIC ACID | 4.6444 | 0.0029448 | 2.5309 | 0.00451520 |
| N-ACETYL-DL-METHIONINE | 4.6358 | 0.0029767 | 2.5263 | 0.00451520 |
| ALPHA-D-GLUCOSE 1-PHOSPHATE | 4.5955 | 0.0031311 | 2.5043 | 0.00469660 |
| L-METHIONINE | 4.3213 | 0.0044328 | 2.3533 | 0.00657610 |
| 3-HYDROXYBUTANOIC ACID | 4.0014 | 0.0067064 | 2.1735 | 0.00984100 |
| 4-GUANIDINOBUTANOATE | 3.6778 | 0.010288 | 1.9877 | 0.01493400 |
| FERULATE | 3.5359 | 0.012445 | 1.9050 | 0.01787400 |
| INDOLE-3-ACETIC ACID | 3.4546 | 0.013892 | 1.8572 | 0.01974100 |
| AZELAIC ACID | 3.4000 | 0.014960 | 1.8251 | 0.02103800 |
| TRANS-CINNAMALDEHYDE | 3.1908 | 0.019919 | 1.7007 | 0.02772200 |
| N-AMIDINO-L-ASPARTATE | 3.0241 | 0.025083 | 1.6006 | 0.03455300 |
| L-TRYPTOPHAN | 2.8945 | 0.030048 | 1.5222 | 0.04097500 |

**Supplementary Table 5: Metabolites with statistically significant differences detected between Y and GF gnotobiotic groups.** List of 22 metabolites with statistically significant differences in relative abundance between Y and GF groups. Statistical significant differences determined by Two-Sided T-test and FDR corrected; P<0.05.

| Compound | T-Stat | P-Value | -Log10(P) | FDR |
|---|---|---|---|---|
| Nicotinic acid | -25.689 | 2.29E-07 | 6.6395 | 2.98E-05 |
| 3-Hydroxybutyric acid | -8.1624 | 0.000182 | 3.7400 | 0.011828 |
| Fumaric acid | -6.6632 | 0.000553 | 3.2575 | 0.023949 |
| L-Asparagine | 5.8051 | 0.001146 | 2.9408 | 0.032220 |
| Ketoleucine | 5.6874 | 0.001275 | 2.8946 | 0.032220 |
| Indoleacetic acid | -5.5199 | 0.001487 | 2.8277 | 0.032220 |
| Acetylalanine | 5.3291 | 0.001780 | 2.7496 | 0.033052 |
| Thymine | -5.0012 | 0.002449 | 2.6109 | 0.037887 |
| Carnosine | 4.9327 | 0.002623 | 2.5812 | 0.037887 |
| 6-Hydroxynicotinic acid | 4.7618 | 0.003121 | 2.5058 | 0.039944 |
| Succinic acid | -4.6555 | 0.003484 | 2.4580 | 0.039944 |
| L-Glutamic acid | 4.5445 | 0.003915 | 2.4073 | 0.039944 |
| Uridine | 4.4241 | 0.004451 | 2.3515 | 0.039944 |
| N-Acetyl-D-tryptophan | 4.4194 | 0.004474 | 2.3493 | 0.039944 |
| L-Serine | 4.3764 | 0.004686 | 2.3292 | 0.039944 |
| Urocanic acid | 4.3323 | 0.004916 | 2.3084 | 0.039944 |
| Docosahexaenoic acid | 4.1881 | 0.005761 | 2.2395 | 0.042696 |
| Shikimic acid | 4.1649 | 0.005912 | 2.2283 | 0.042696 |
| Suberic acid | 4.0754 | 0.006536 | 2.1847 | 0.044720 |
| Citraconic acid | 4.0233 | 0.006933 | 2.1591 | 0.045064 |
| L-Cystathionine | -3.9504 | 0.007534 | 2.1230 | 0.046639 |
| N-Acetyl-L-phenylalanine | 3.8837 | 0.008136 | 2.0896 | 0.048077 |
